# Dissociable neural systems for unconditioned acute and sustained fear

**DOI:** 10.1101/676650

**Authors:** Matthew Hudson, Kerttu Seppälä, Vesa Putkinen, Lihua Sun, Enrico Glerean, Tomi Karjalainen, Jussi Hirvonen, Lauri Nummenmaa

**Affiliations:** Turku PET Centre, University of Turku, Turku, Finland; Department of Neuroscience and Biomedical Engineering, School of Science, Aalto University, Espoo, Finland; Department of Radiology, Turku University Hospital, Turku, Finland; Department of Psychology, University of Turku, Turku, Finland

**Keywords:** Fear, Threat, Neural Synchronization, Horror Movies, Naturalistic fMRI

## Abstract

Fear protects organisms by increasing vigilance and preparedness, and by coordinating survival responses during life-threatening encounters. The fear circuit must thus operate on multiple timescales ranging from preparatory sustained alertness to acute fight-or-flight responses. Here we studied the brain basis of sustained (“looming”) and acute fear using naturalistic functional magnetic resonance imaging (fMRI) enabling analysis of different time-scales of fear responses. Subjects (N = 51) watched feature-length horror movies while their hemodynamic brain activity was measured with fMRI. Time-variable intersubject correlation (ISC) was used to quantify the reliability of brain activity across participants, and seed-based phase synchronization was used for characterizing dynamic connectivity. Subjective ratings of fear were obtained from a separate sample, and were used to assess how synchronization and functional connectivity varied with emotional intensity. These data suggest that acute and sustained fear are supported by distinct neural pathways, with sustained fear amplifying mainly sensory responses, and acute fear increasing activity in brainstem, thalamus, amygdala and cingulate cortices. Sustained fear increased ISC in regions associated with acute fear, and also amplified functional connectivity within this network. The results were replicated in two independent experiments with different subject samples. The functional interplay between cortical networks involved in sustained anticipation of, and acute response to, threat involves a complex and dynamic interaction that depends on the proximity of threat, and the need to employ threat appraisals and vigilance for decision making and response selection.

## Dissociable neural systems for unconditioned acute and sustained fear

Emotions prepare us for action. They motivate seeking out rewarding stimuli, increasing alertness and avoiding threat (Bernhardt & Singer, 2012). Fear has a strong developmental and evolutionary function as a primordial reaction to danger that elicits a distinctive physiological and psychological response. The endocrine system releases epinephrine, norepinephrine, and cortisol that excites the cardiovascular and respiratory systems, and releases glucose into the bloodstream, preparing the body for physical action (Rodrigues, LeDoux, & Sapolsky, 2009). A concomitant increase in attentional vigilance (Finucane, 2011) and a bias toward threatening stimuli (Öhman, Flykt, & Esteves, 2001) serve to heighten perceptual awareness, and learning/memory mechanisms (Öhman & Mineka, 2001). Fear is associated with changes in both the central and peripheral nervous system (Ekman, 1992; Kreibig, 2010; Nummenmaa & Saarimäki, 2017; Panksepp, 1982) and is also characterized by an idiosyncratic subjective experience (Nummenmaa, Glerean, Hari, & Hietanen, 2014; Nummenmaa, Hari, Hietanen, & Glerean, 2018) and overt expression (Smith, Cottrell, Gosselin, & Schyns, 2005).

Acute fear is associated with a distinctive pattern of neural activity distributed through the cerebellum (Ploghaus et al., 1999), limbic system (Knight, Smith, Cheng, Stein, & Helmstetter, 2004; LaBar, Gatenby, Gore, LeDoux, & Phelps, 1998), and cortex (*prefrontal:* Phelps, Delgado, Nearing, & LeDoux, 2004; *sensory:* Morris, Buchel, & Dolan, 2001; *cingulate:* Milad, Quirk, Pitman, Orr, Fischl, & Rauch, 2007; *insula:* Critchley, Mathias, & Dolan, 2002; *motor:* Lissek et al., 2014). This distributed network (Saarimäki et al., 2016) enables the rapid detection of potential threat and its precursors, the appraisal of the threat and its saliency to oneself, the employment of executive functioning and memory for decision making and action planning, and the implementation of action plans (Zhu & Thagard, 2002).

In addition to generating immediate survival responses, fear systems also modulate vigilance in anticipation of threat caused by environmental cues, perceptual uncertainty, and ambiguity (Fanselow, 1994; Lang, Davis, & Öhman, 2000; Lehne & Koelsch, 2015). This gives rise to subjective feelings of anxiety, tension, suspense, dread, or foreboding that reflects a generalized anticipatory preparedness for the possibility of potential danger. Several recent studies have shown that spatiotemporally distant threats elicit activity in the ventromedial prefrontal cortex, posterior cingulate cortex, hippocampus and amygdala, which are associated with a cognitive mechanism of fear that reflects the need for complex information processing and memory retrieval to generate an adaptive and flexible response. A threat that is proximal in space or time, on the other hand, elicits a reactive fear response of immediate action and fight or flight, and which elicits activity in the periaqueductal gray, amygdala, hypothalamus, and middle cingulate cortex (Mobbs et al., 2007; Qi, Hassabis, Sun, Guo, Daw, & Mobbs, 2018).

To date, studies on the differential anticipatory and reactionary networks of fear have compared events that predict and follow the onset of threat, but employed a discrete and categorical distinction that collapses the temporal dimension, thus failing to capture the temporal dynamics and fluctuation of the fear response. Therefore, the neural mechanisms supporting dynamically fluctuating and sustained fear, versus acute fear responses, during naturalistic conditions remains poorly characterized. Furthermore, the majority of human neuroimaging studies have been conducted using relatively impoverished and reduced laboratory settings, which does not necessarily provide an optimal model of how the brain responds to survival challenges in the real world (see review in Adolphs, Nummenmaa, Todorov, & Haxby, 2016). First, the brain has evolved to parse the dynamic world and its complex events, and it is known that neural responses to complex stimuli cannot necessarily be predicted on the statistical combination of responses to simple stimuli (Felsen & Dan, 2005). Cells in the cat visual cortex show stronger responses to real pictures than to random patterns (Touryan, Felsen, & Dan, 2005), and gamma band responses and local field potentials are also most reliable in response to repeated presentations of movies (Belitski et al., 2008). In humans, life-like moving faces also elicit markedly stronger activation in the face processing network than static or rigidly moving faces (Fox, Iaria, & Barton, 2009; Schultz, Brockhaus, Bulthoff, & Pilz, 2013). And most importantly, many psychological phenomena – including fear – span multiple overlapping time scales and parallel processing of multiple features, thus they cannot be adequately studied with rigid and tightly controlled classical experimental designs.

Recent advances in brain signal analysis have however enabled characterization of brain activity during naturalistic and unconstrained conditions where the stimulus space is high-dimensional (Nummenmaa, Lahnakoski, & Glerean, 2018). During natural audiovisual stimulation, subjects’ brain activation becomes synchronized in occipital and temporal regions of the cortex, due to the identical perceptual experience of the participants (Glerean, Salmi, Lahnakoski, Jääskeläinen, & Sams, 2012; Hasson, Nir, Levy, Fuhrmann, & Malach, 2004). The spatial distribution of the synchronization is functionally specific: for example, greater synchronization in the fusiform gyrus is found during portions of the movie when faces are visible (Hasson, Furman, Clark, Dudai, & Davachi, 2008). Such neural synchronization is subject not only to bottom-up changes in perceptual input but top-down changes in attentional control (Lahnakoski et al., 2014). Importantly, activity in regions involved in the perception and experience of emotions become increasingly synchronized across individuals as a function of the emotional content of the stimulus (Nummenmaa, Glerean, Viinikainen, Jääskeläinen, Hari, & Sams, 2012; Nummenmaa, Saarimäki, Glerean, Gotsopoulos, Hari, & Sams, 2014). For example, changes of emotional valence from positive to negative alters synchronization in regions such as the thalamus, anterior cingulate cortex, and prefrontal cortex. In contrast, the arousal elicited by an emotional stimulus alters synchronization in visual and somatosensory regions. These methods can capture the complex temporal dynamics of neural activity in response to naturalistic stimuli for which controlled modelling is not possible, but nevertheless reveal reliable neural activity at the population level on a moment-to-moment basis in functionally specific brain regions.

Additionally, the prolonged and variable brain activation time series resulting from naturalistic stimulation is well-suited for analyzing dynamic connectivity changes (Nummenmaa, Saarimäki, et al., 2014). Prior studies on emotion-dependent brain connectivity have typically constrained the analysis to a small numbers of regions of interest and suffered from the low power of conventional event-related and boxcar designs in connectivity analyses (e.g. Eryilmaz, Van De Ville, Schwartz, & Vuilleumier, 2011; Passamonti, Rowe, Ewbank, Hampshire, Keane, & Calder, 2008; Tettamanti, Rognoni, Cafiero, Costa, Galati, & Perani, 2012). A naturalistic stimulation setup in turn offers a high-powered alternative for tapping fear-dependent connectivity in the brain.

## The present study

The aim of the current study was to investigate, in naturalistic settings, the neural mechanisms involved in generating acute fear responses and those supporting sustained anticipatory fear when the threat is not yet present. Subjects viewed feature-length horror movies while their brain activity was recorded with fMRI. Acute threatening events (“jump scares”) were annotated in the movies, and self-reports of sustained fear were obtained. These time series were subsequently used for predicting hemodynamic activity, voxel-wise intersubject correlation, and functional connectivity. We show that acute and sustained fear are supported by distinct neural pathways. Importantly, we confirm the consistency of these effects using a voxel-wise intraclass-correlation reliability measure across two independent data sets with different subjects and stimuli.

## Materials and methods

The study protocol was approved by the ethics board of the hospital district of Southwest Finland, and the study was conducted in accordance with the Declaration of Helsinki.

### Stimuli

Two feature length horror movies (The Conjuring 2, 2016, and Insidious, 2010, both directed by James Wan) were selected based on a pilot survey on horror movies. Online databases (Rotten Tomatoes, International Movie Database, AllMovie) were consulted to generate a list of the 100 best-rated horror movies of the past 100 years. An independent sample of 216 participants completed a survey asking if they had seen the movies and, if so, rated them on scariness and quality. The number of jump scares in each movie were indexed from an online database (http://wheresthejump.com, 2017). The participants also reported how often they watched horror movies or movies in general, and how scary they considered different types of horror (e.g. psychological horror, supernatural horror). Finally, the participants reported the most common emotions experienced while viewing horror movies. These data (**Figure 1**) confirmed that viewing horror movies was common, a total of 72% of respondents reported watching at least one horror movie every six months. Psychological horror and movies based on supernatural as well as real events were rated as most frightening and viewing horror movies was associated with the targeted emotions (fear, anxiety, excitement). Data for the exposure, and fear and quality ratings for the top 10 scariest movies are shown in **Table 1** (see **Supplementary Analysis 2.** for the full results). Scariness and quality ratings (Bonferroni corrected alpha = 0.017) were positively correlated (*r* = .528, *p* < .001) showing that movies considered as high quality were also considered as scary.

**Figure 1.**
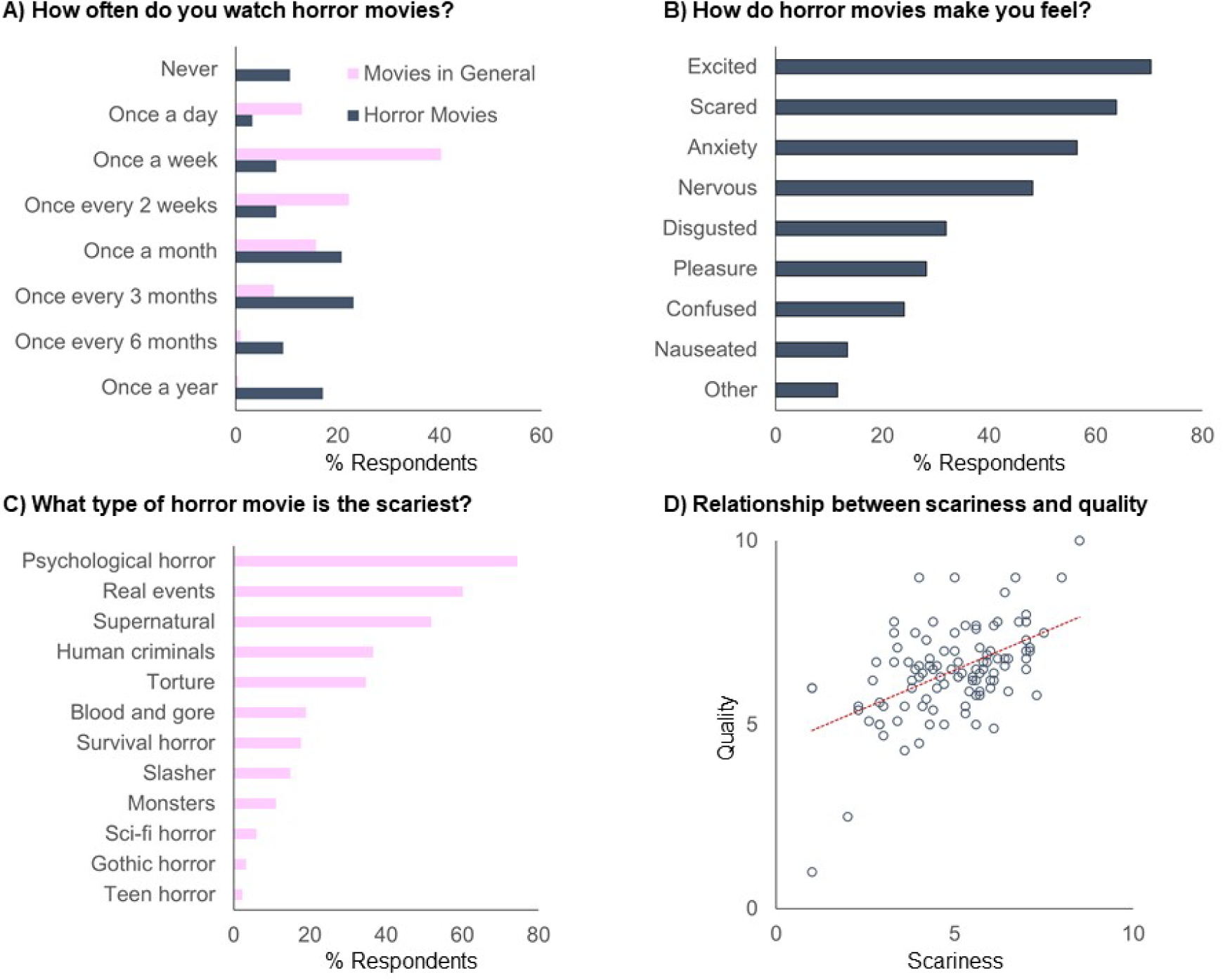
Emotional responses to horror movies. The frequency of viewing horror movies by the sample (A), and the different emotions experienced while viewing horror movies (B), as well as the type of horror movie that participants found to be the scariest (C). Lastly, the relationship between the scariness of the horror movie and its quality (D).

**Table 1.** Top 10 scariest movies in the survey. Title and year of release, alongside familiarity with the film, and scariness and quality ratings by those who had seen it, and also the number of jump scares (where available). Selected movies indicated in bold.

| Title | Year | Have you seen this movie? |  |  | Scary | Quality | Jump scares |
| --- | --- | --- | --- | --- | --- | --- | --- |
|  |  | Yes | No, but heard of it | Not seen nor heard of it |  |  |  |
| The Devil's Backbone | 2001 | 4.2 | 6.3 | 89.6 | 8.5 | 10.0 | 4 |
| The Wailing | 2016 | 2.1 | 8.3 | 89.6 | 8.0 | 9.0 | 4 |
| The Conjuring | 2013 | 39.6 | 31.3 | 29.2 | 7.5 | 7.5 | 12 |
| REC 2 | 2009 | 10.4 | 29.2 | 60.4 | 7.3 | 5.8 | - |
| **Insidious** | **2010** | **29.2** | **25.6** | **45.2** | **7.1** | **7.1** | **24** |
| The Exorcist | 1973 | 53.0 | 23.2 | 23.8 | 7.1 | 7.0 | 10 |
| Goodnight Mommy | 2015 | 4.2 | 10.4 | 85.4 | 7.0 | 8.0 | 1 |
| A Chinese Ghost Story | 1987 | 1.8 | 7.1 | 91.1 | 7.0 | 7.8 | - |
| **The Conjuring 2** | **2016** | **37.5** | **31.0** | **31.5** | **7.0** | **7.3** | **22** |
| Under The Shadow | 2016 | 2.1 | 14.6 | 83.3 | 7.0 | 7.0 | 9 |

The selected movies were chosen on the basis of this survey as 1) they were rated highly for scariness (Conjuring 2 = 7.0; Insidious = 7.1) and quality (Conjuring 2 = 7.3; Insidious = 7.1), 2) they contained frequent jump scare events (Conjuring 2 n = 22; Insidious n = 24), and 3) relatively few people had seen them (Conjuring 2 = 17.4%; Insidious = 13.5%). Each movie was edited for length with Apple iMovie whilst maintaining the fear elements of the film (durations: The Conjuring 2 = 109 minutes, Insidious = 94 minutes). The movies were split into short segments of approximately 13 minutes (9 for The Conjuring 2, 7 for Insidious) to optimize stimulus delivery and data processing, and to allow short breaks for the subjects. Cut-points adhered to naturally occurring scene transitions within the films. The videos were presented at 23.98fps at a resolution of 800 * 600.

### Participants

A total of fifty-one subjects took part in the brain imaging and behavioral experiments (mean age = 24.5, SD = 5.5, 30 females). Imaging data were obtained from thirty-seven participants (Conjuring 2: n = 18, Insidious: n = 19). One further subject was scanned (The Conjuring 2) but the scan had to be terminated due to subject discomfort. Behavioral fear ratings were obtained from 40 subjects (Conjuring 2 n = 20, Insidious n = 20, an additional subject who rated both movies was excluded for not complying to task instructions). Twenty-eight subjects viewed both movies, although no subject viewed the same movie twice (See **Table 2**. for details). Subjects were recruited from the University of Turku and surrounding community, they received payment and/or course credit as compensation, and provided written informed consent prior to taking part. No participants had a history of neuropsychiatric symptoms or medication.

**Table 2.** Distribution of subjects for whom imaging data and/or behavioral fear ratings were acquired across the two movies (prior to subject exclusions).

| Movie |  | The Conjuring 2 |  |  |
| --- | --- | --- | --- | --- |
|  |  | Scan | Rate | Not Viewed |
| Insidious | Scan | 1 | 14 | 4 |
|  | Rate | 11 | 3 | 7 |
|  | Not Viewed | 7 | 4 | - |

### Procedure

#### Behavioral Fear Ratings

Subjects viewed the movie on an iMac retina 5K 27-inch monitor running High Sierra 10.13.3. Audio was delivered via AKG K550 MKII headphones (32 ohms, 114db, 12hz to 28kHz). A vertically oriented slider ranging from 0 (no fear) to 1 (maximum fear) was placed to the right of the movie window. Participants controlled the location of a cursor on the scale by moving it upwards (push the mouse forward) when their fear increased and moving the cursor downwards (pull the mouse backward) when their fear decreased. The cursor position was monitored at 5Hz. Subjects were told to use the full range of the scale and to make sure that their rating continuously reflected how scared they were. Stimuli were presented in similar segments as described above. A timer was placed inconspicuously at the top of the screen that counted down until the end of the current segment. Participants were told that they were free to take a break but that they should wait until the end of the current segment before they did so.

#### fMRI Acquisition and Pre-processing

MR imaging was conducted at Turku PET Centre at the University of Turku using a Philips Ingenuity TF 3-Tesla scanner. Anatomical images (1 mm^3^ resolution) were acquired using a T1-weighted sequence (1mm^3^ resolution, TR 8.1ms, TE 3.7ms, flip angle 7°, 256mm FOV, 256 × 256 acquisition matrix, 176 sagittal slices). Whole-brain functional data were acquired with T2*-weighted echo-planar imaging sequence, sensitive to the blood-oxygen-level-dependent (BOLD) signal contrast (TR = 2600ms, TE = 30ms, 75° flip angle, 250 mm FOV, 80 × 80 acquisition matrix, 8.132/53.4Hz bandwidth, 3.0 mm slice thickness, 45 interleaved slices acquired in ascending order without gaps).

Scanning was split into short runs of approximately 25 minutes each (comprising 2 to 3 segments) for the sake of subject comfort (number of volumes: The Conjuring 2, 675, 696, 595, and 572 scans (total 2538 scans); Insidious: 725, 780, and 687 scans (total 2192 scans)) with breaks between each run. Stimulus video was displayed using goggles affixed to the head coil (NordicNeuroLab VisualSystem). Audio was played through SensiMetrics S14 earphones (100Hz – 8 kHz bandwidth, 110dB SPL). Volume was adjusted to a comfortable level that could still be heard over the scanner noise.

Functional data were preprocessed with FSL (www.fsl.fmrib.ox.ac.uk) using the FEAT pipeline. The EPI images were realigned to the middle scan by rigid body transformations to correct for head movements (six parameters). Two-step coregistration was conducted firstly to the participant’s T1 weighted structural image, and secondly to MNI152 2mm template. Spatial smoothing used isotropic Gaussian kernel whose full width at half maximum (FWHM) was 8 mm. Low-frequency drifts in data were estimated and removed using a 240s Savitzky-Golay filter (Cukur, Nishimoto, Huth, & Gallant, 2013).

### Data Analysis

#### General Linear Model Analysis

GLM analyses were implemented with SPM 12 (www.fil.ion.ucl.ac.uk/spm) with a two-stage random effects analysis. All results were thresholded with a cluster-level FDR corrected alpha of 0.001, unless noted otherwise, and an extent threshold of 10 voxels. Frame-wise luminance and sound intensity time series were derived using custom Matlab toolbox and down-sampled to 2.6 seconds (1TR). These were entered as nuisance regressors in all analyses; however, they did not alter the results appreciably. All results are therefore reported without these regressors. Primary second-level analyses were conducted together for both movies, with separate analyses reported in the **Supplementary Materials**.

#### Jump-scares

We first modelled the brain responses to acute fear. In the first-level analysis we modelled jump-scares with stick functions convolved with a canonical hemodynamic response function. High-pass filter period was set to 128s. Second-level analysis was conducted on the resulting contrast images using a one sample t-test.

#### Dynamic Fear Ratings

Sustained fear responses were analyzed using the behavioral fear ratings averaged across subjects. In the first-level analysis, the TR down-sampled fear ratings were convolved with a canonical hemodynamic response function and entered as a regressor into the GLM analysis (high-pass filter period 256s). Second-level analysis on these contrasts was conducted using a one sample t-test.

#### Intraclass correlation

To provide a statistical estimate of reliability of the observed effects, intraclass correlation (ICC) analyses were conducted across the two samples (see Bennett & Miller, 2010; Chen et al., 2017). A one-way random effects between-subjects ICC (1,1) (Shrout & Fleiss, 1979) was conducted for each voxel. The t-contrast for each participant at the first level of analysis was subject to a one-way between subjects ANOVA, and the between subjects (BS) and within subjects (WS) mean square (MS) used to compute the ICC by dividing the target variance (BS_MS – WS_MS) over the total variance (BS_MS + (2*WS_MS)). The variance ratio varies between 0 and 1, with higher values reflecting greater reliability (Koo & Li, 2016). The ICC maps for the jump-scares and dynamic fear ratings are presented in the **Supplementary Materials**.

#### Inter-Subject Correlation Analysis (ISC)

ISC provides a model-free means for analyzing hemodynamic responses to complex naturalistic stimuli (Hasson et al., 2004). ISC was implemented using the ISC toolbox (release 21: https://www.nitrc.org/projects/isc-toolbox/). Detailed information can be found at Kauppi, Pajula & Tohka (2014). Pearson correlation coefficient for each subject pair voxel time series provided a measure of across subject similarity in BOLD activity.

ISC was calculated in two ways. First, mean ISC was computed for the full time series to provide an average ISC map during the course of the whole movie. This reveals the typical time-locking of neural responses across subjects throughout the movie. Statistical significance of the ISC values was calculated by means of a fully nonparametric voxel wise permutation test of the r value. Each subject’s time series was circularly shifted by a random number of time points, which preserved the temporal autocorrelations present within each time series but disrupted the temporal alignment between subjects. The ISC value was computed over 1,000,000 realizations, and the resulting distribution of r values represent the non-correlated time series that would be expected if the null hypothesis were true. The *p* values were FDR thresholded at p = 0.05 without assumptions.

Second, a dynamic measure of neural synchronization was calculated to establish how intersubject synchronization varies throughout the movie. Such a dynamic approach allows modelling of whether moment-to-moment ISC is associated with the currently experienced fear level of the participants. Voxel-wise ISC values were computed for each time point with a sliding window of 10 samples (see Nummenmaa et al., 2012). Dynamic ISC time series were computed separately for each session. As each session lost 9 time points due to moving averaging, the voxel time-series were de-meaned and the sliding window ISC was calculated for the concatenated final 9 volumes of one session and the first 9 volumes of the next session, producing the ISC for the missing 9 time points. This provided a continuous ISC measure across the whole movie (minus the final 9 samples). Fear ratings (down-sampled to 1TR) were also subjected to similar moving averaging with a 10-sample sliding window to match their temporal resolution with the dynamic ISC signal. Pearson correlation coefficient between the ISC time series and the fear rating were then computed to reveal how across subject similarity in neural activity is associated with feelings of fear. The assumption of independence between data points in the voxel time series and the fear ratings were not met, therefore p values were calculated with a corrected degrees of freedom to account for autocorrelations in the data (Conjuring 2: 103 to 690; Insidious: 95 to 539; see Alluri, Toiviainen, Jääskeläinen, Glerean, Sams, & Brattico, 2012; Pyper & Peterman, 1998), with a FDR thresholded alpha of 0.05. Note that acute fear events could not be meaningfully analyzed with this technique, as predicting the moving-averaged ISC time series with stick functions with zero duration would not be conceptually meaningful.

### Functional Connectivity Analysis

Functional Connectivity was investigated by employing Seed Based Phase Synchronization (SBPS), implemented with the FunPsy toolbox (https://github.com/eglerean/funpsy) and described in detail in Glerean et al., (2012). For each participant, the demeaned BOLD time series for each voxel was band-pass filtered to remove cardiovascular noise (0.04 – 0.07 Hz). Regions of interest (45 per hemisphere, 26 cerebellar) were defined based on the AAL atlas in MNI space (Tzourio-Mazoyer et al., 2002) and the BOLD signal was averaged across voxels in each region. The phase analytic signal (in radians) of the Hilbert transformed BOLD response of each region was calculated. The instantaneous angular difference between each region pair at each time point provides a model free estimation of dynamic neural synchronization between regions that maintains the temporal precision that is lost when using a sliding window analysis. The time series of neural synchronization between each pair of regions was then correlated with the fear ratings to assess how functional connectivity altered with the fear of the participant. Alpha levels were subject to FDR correction using corrected degrees of freedom based on the autocorrelation between the angular difference and fear ratings (Conjuring 2: 171 to 753; Insidious: 100 to 598).

## Results

### Behavioral Data

Behavioral data (**Figure 1**) revealed that both movies elicited strong subjective feelings of fear that were also varying over time. Ratings were consistent across subjects, as indicated by the high ISC of the fear time series (mean z transformed Pearson’s *r*: Conjuring 2 = .756; Insidious = .630) and low mean 95% CI (Conjuring 2 = .08; Insidious =.09). Accordingly, averaged ratings provide a good model for experienced fear during the fMRI experiment. Interestingly, there were significant positive correlations between fear ratings and 95% CI and ISC for both movies (all p’s < .001), suggesting that, as the intensity of fear increased, variability in *absolute* ratings of fear increased, but the extent to which participant’s ratings followed each other (synchronized) also increased. In other words, during fearful events participants’ experience of fear showed a general increase of different orders of magnitude, which also led to more similar time-locking of subjective feelings across participants. Peaks of the self-reported fear time series were concordant with the occurrence of jump-scares (black vertical lines in **Figure 1**). The fear ratings correlated negatively with luminance (The Conjuring 2: r = -.433, p < .001; Insidious: r = -.253, p <.001), and positively with sound intensity (The Conjuring 2: r = .408, p < .001; Insidious: r = .368, p <.001).

**Figure 1.**
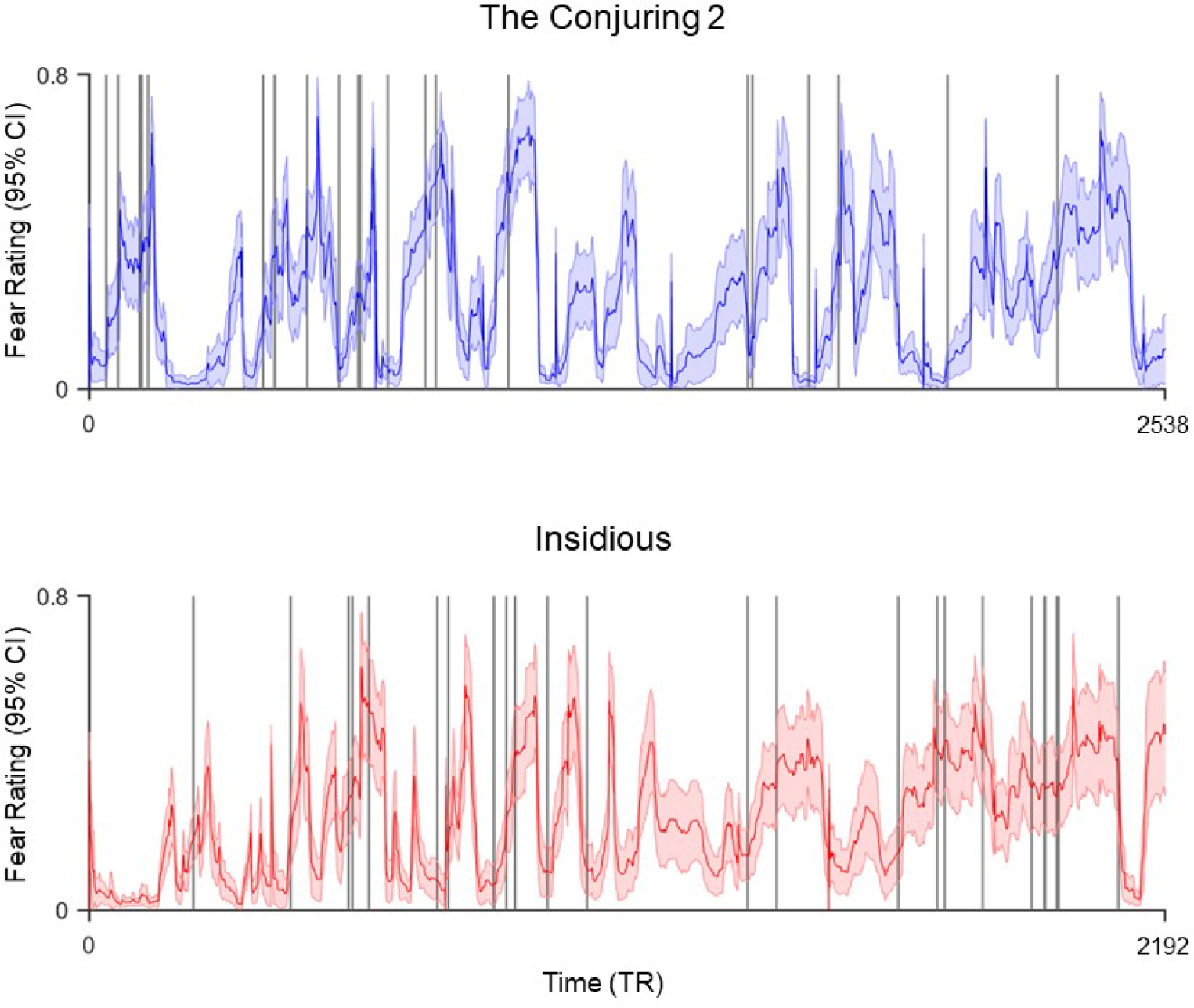
Mean dynamic fear ratings (scaled from 0 to 1) with 95% confidence intervals. Black vertical lines represent the jump scare events (Conjuring 2 n = 22; Insidious n = 24).

### The effect of acute transient fear on neural activity

BOLD responses to jump-scare events (joint analysis of both movies) are shown in **Figure 2**. The mean ICC coefficient across the two movies was 0.65 (indicative of moderate reliability, Koo & li, 2016). There was extensive bilateral activity in the cuneus, precuneus, lingual gyri, middle occipital gyri, and fusiform gyri. Parietal activity was observed in bilateral precentral gyrus. In the temporal lobe, bilateral activity was observed in the posterior, middle, and anterior portions of the superior temporal gyri, as well as the middle and transverse temporal gyri. Frontal activity was observed in bilateral posterior and anterior portions of the inferior frontal gyrus, and medial frontal gyrus, and the cingulate cortex also exhibited robust activity in the anterior, middle and posterior portions. Bilateral insula cortex activity was evident in anterior and posterior regions, and also the claustrum. There was bilateral amygdala and parrahippocampul activity, as well as subcortical activity in the caudate, thalamus, putamen, and the red nucleus and substania nigra of the periaqueductal gray area. A large cluster of activity was also found in the cerebellum, encompassing the cerebellar tonsil, culmen, declive, pyramis, nodule, uvula, fastigium, and cerebellar lingual. No regions showed a decrease in activity in response to the jump-scares after controlling for the false discovery rate. However, with an uncorrected threshold of *p* < .001, small bilateral clusters in the posterior anterior cingulate cortex, parrahippocampus, and caudate were evident.

**Figure 2.**
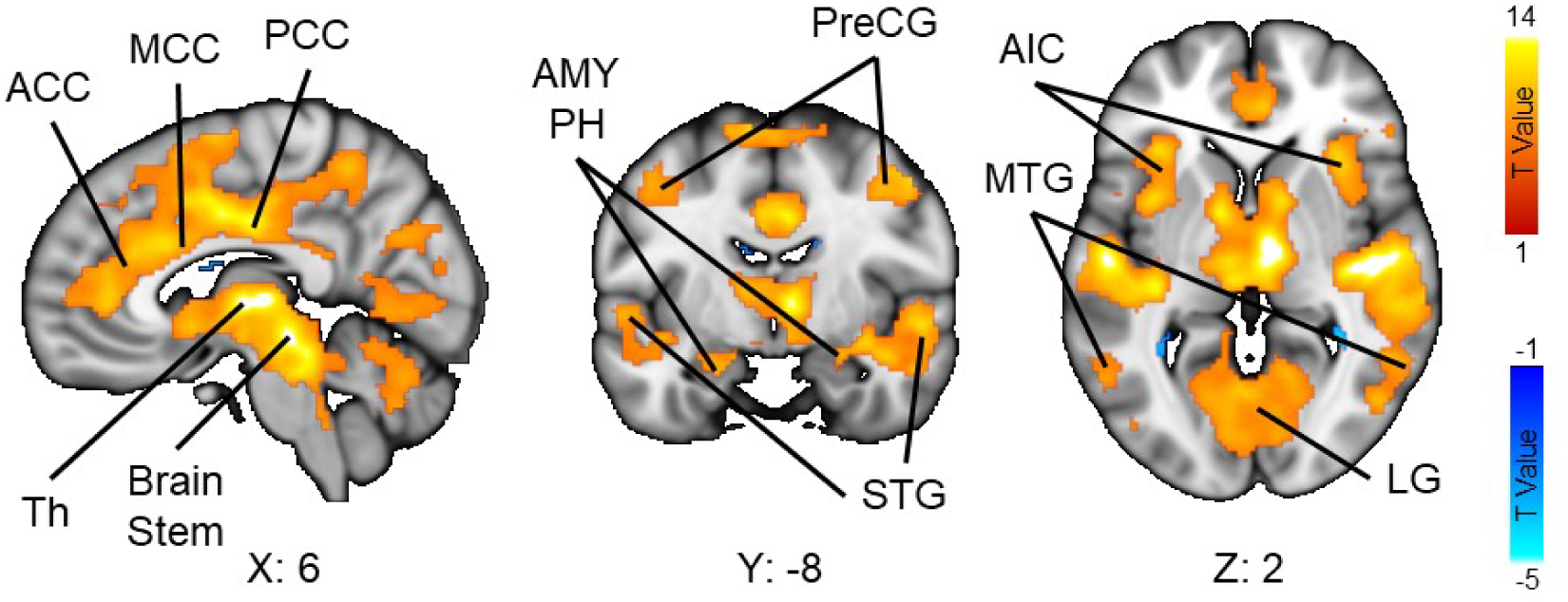
The effect of acute fear (jump scares) on neural activity collapsed across movies (FDR corrected p = 0.001). ACC: Anterior Cingulate Cortex, MCC: Middle Cingulate Cortex, PCC = Posterior Cingulate Cortex, Th = Thalamus, AMY = Amygdala, PH = Parrahippocampus, PreCG = PreCentral Gyrus, STG = Superior Temporal Gyrus, AIC = anterior Insula Cortex, MTG = Middle Temporal Gyrus, LG = Lingual Gyrus. Results for each movie and the intraclass correlation analysis are in **Supplementary Figure 1.**

### The effect of sustained fear on neural activity

Regions that exhibited activity significantly correlated with the sustained level of fear are shown in **Figure 3**. The mean ICC coefficient across movies was 0.61 (indicative of moderate reliability, Koo & Li, 2016). Fear predicted activity in bilateral posterior middle occipital gyri, left fusiform gyri, and cuneus, and also the right lingual gyrus and right precuneus. Cerebellar activity was evident in left uvula, and bilateral declive, culmen, and pyramis. No regions exhibited a negative relationship with fear at a FDR corrected threshold of p <. 001. However, at a FDR corrected threshold of p < .05, decreased activity was observed in bilateral post-central gyrus, and bilateral inferior parietal lobe extending to the supramarginal gyrus in the left hemisphere. In the frontal lobe, there was activity in the left ventral and dorsal inferior frontal gyri, left ventral medial frontal gyrus, left ventral and dorsal middle frontal gyrus, left precentreal gyrus, and bilateral medial frontal gyri. Regions of the insular cortex (left anterior and right middle) also exhibited decreased activity with rising fear, as did the left claustrum, and decreases in activity were also observed in bilateral parrahippocampus, caudate, and thalamus. However, when luminance and sound intensity were added as covariates, a positive relationship with fear was observed only in small clusters in the right parahippocampul gyrus and right lingual gyrus, suggesting that the majority of associated activity with fear ratings was driven by the contribution of stimulus features such as intense sounds and diminished/uncertain visual input. The pattern of negative associations with fear was more stable after accounting for luminance and sound intensity (bilateral posterior superior temporal gyri, left precentral gyrus, left dorsal inferior frontal gyrus, right dorsal superior frontal gyrus, bilateral anterior insula cortex, bilateral anterior cingulate cortex, and caudate).

**Figure 3.**
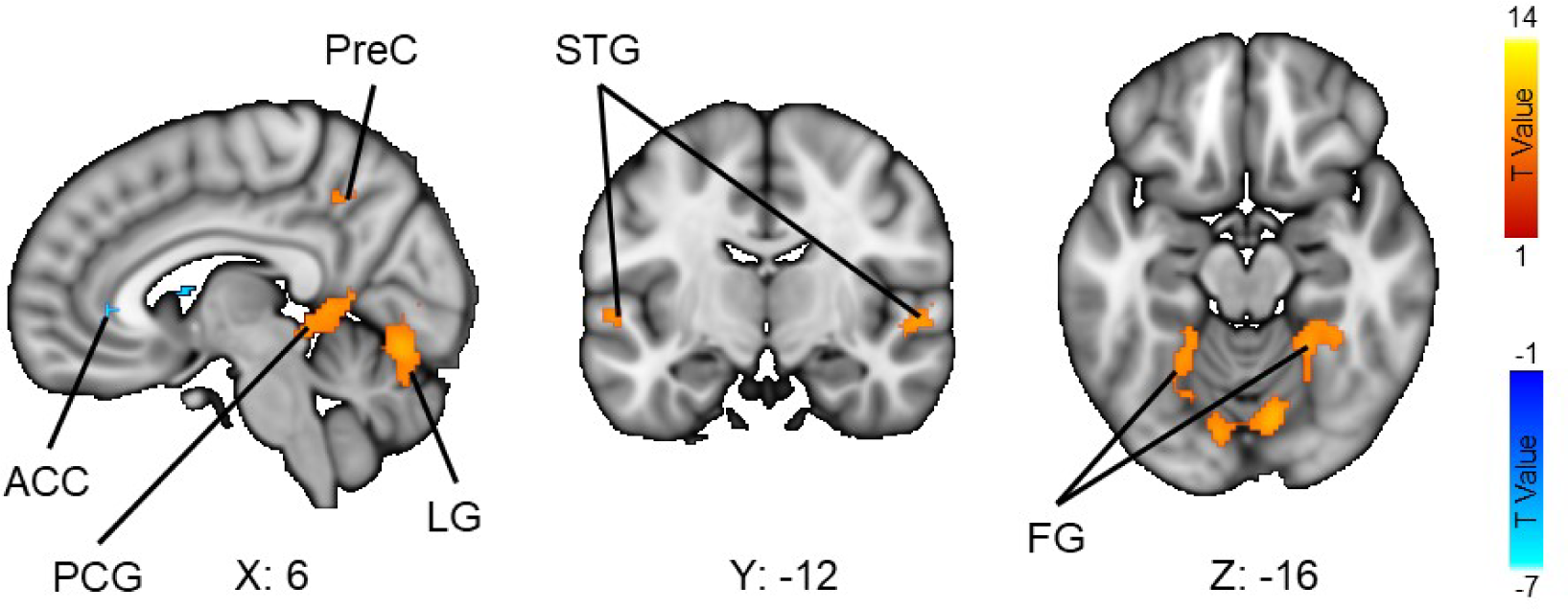
The relationship between sustained fear and neural activity across both movies. (FDR corrected *p* = 0.001). ACC = Anterior Cingulate Cortex, PCG = Post-Cingulate Gyrus, LG = Lingual Gyrus, PreC = Precuneus, STG = Superior Temporal Gyrus, FG = Fusiform Gyrus. Results for each movie and the intraclass correlation analysis are in **Supplementary Figure 2.**

### The effect of sustained fear on intersubject synchronization of brain activity (ISC)

Mean ISC maps for each movie are shown in **Figure 4**. For both movies, synchronized activity was evident across the whole brain, but there was a clear gradient from higher synchronization in the sensory cortices to parietal areas and association cortices, to the lowest synchronization in frontal regions.

**Figure 4.**
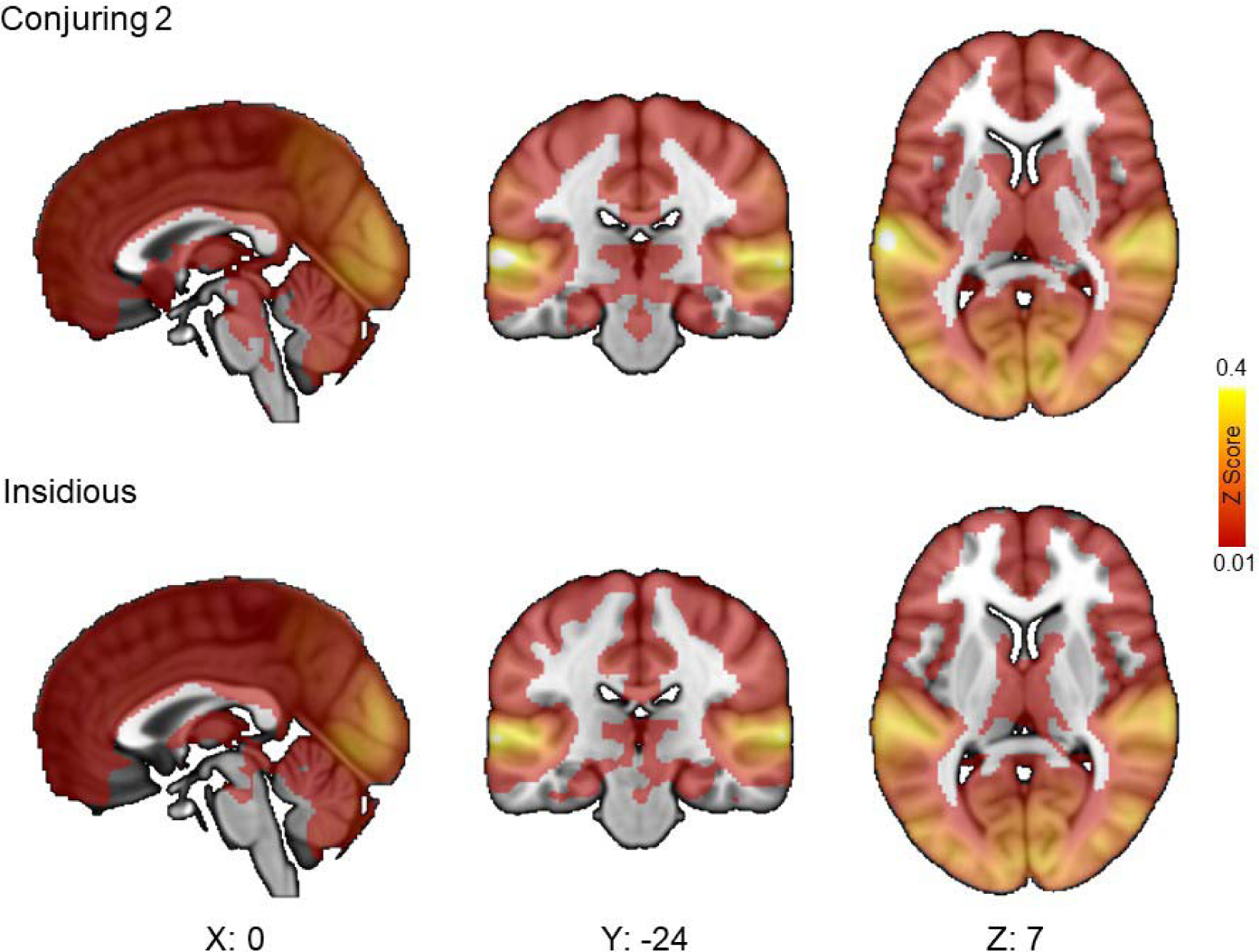
Intersubject Correlation Maps. Voxel intensities show mean z transformed Pearson’s correlation coefficient for each subject pair’s voxel-wise time series across the whole movie. Statistical significance of the ISC values was calculated by means of a fully nonparametric voxel wise permutation test of the r value (1,000,000 realizations, FDR corrected *p* = 0.05).

The relationship between the dynamic ISC and the sustained fear is depicted in **Figure 5**. Intensity of fear was associated with increased ISC as the level of fear rose in a large bilateral swathe of cortex from the anterior and middle portions of the cingulate gyrus, to the medial frontal gyrus and paracentral lobule, to the primary somatosensory cortex (postcentral gyrus) and the adjacent precentral gyri. This effect was evident in both movies. Replicable effects were also found in the left superior frontal gyrus (also right hemisphere for The Conjuring 2), bilateral inferior frontal gyri, bilateral posterior, middle and anterior insula cortices, and bilateral thalamus. Both movies exhibited a fear dependent increase in ISC in posterior nodes of the frontoparietal attention circuits (bilateral inferior parietal cortices) and precuneus, the temporal lobe (right anterior superior temporal gyrus, bilateral middle temporal gyrus), and occipital lobe (left post cingulate gyrus for both movies, right post-cingulate gyrus). The fear related increase in ISC was also evident in the cerebellum for both movies (culmen, bilateral tuber, pyramis, devlice and inferior semi-lunar lobe).

**Figure 5.**
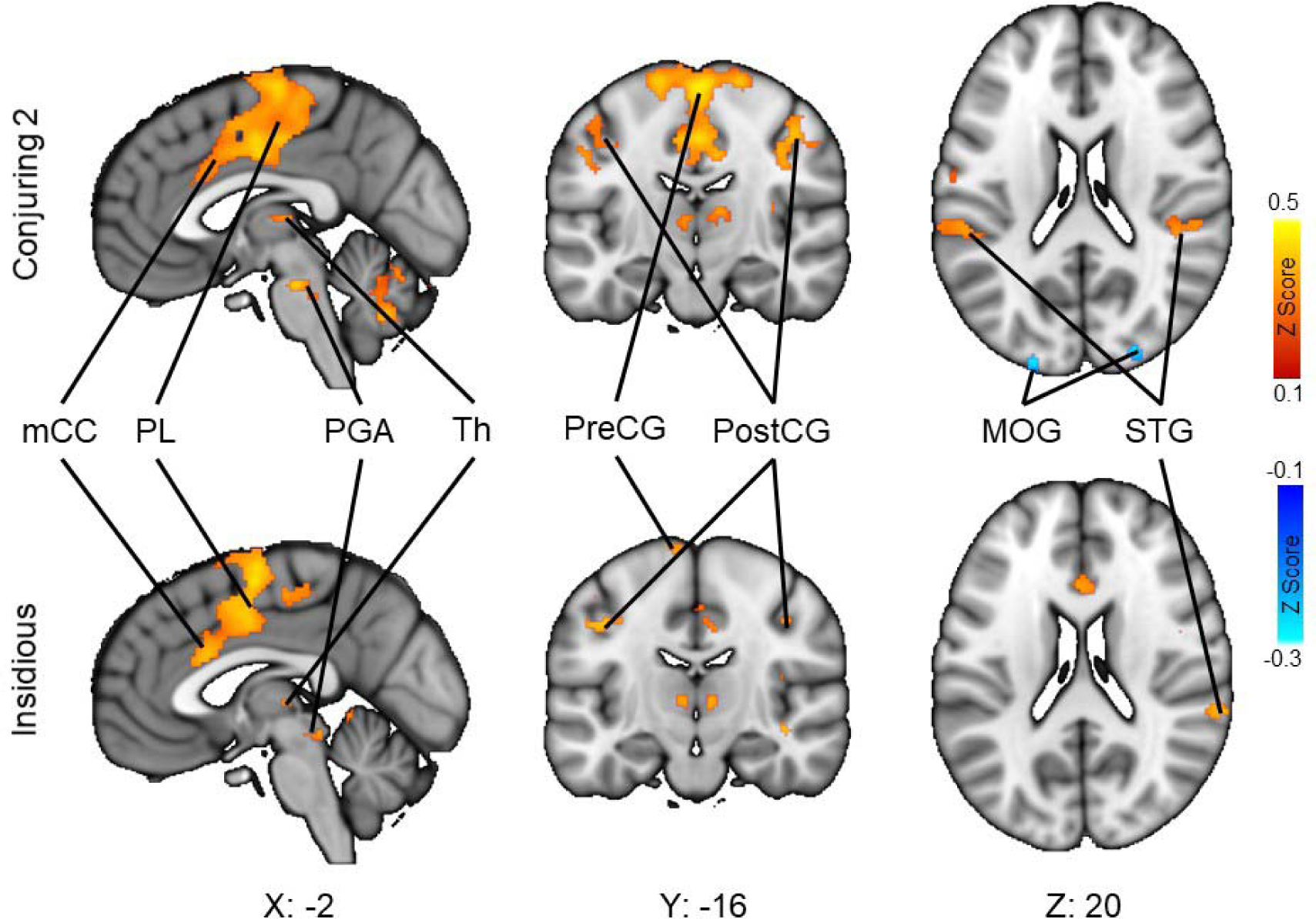
Fear-dependent dynamic inter-subject neural synchronization for The Conjuring 2 (top) and Insidious (bottom). The data are shown as z-transformed Pearson’s r (FDR corrected *p* = .001). mCC = Middle Cingulate Cortex, PL = Paracentral Lobule, PGA = Periaqueductal Gray Area, Th = Thalamus, PreCG = PreCentral Gyrus, Post CG = PostCentral Gyrus, MOG = Middle Occipital Gyrus, STG = Superior Temporal Gyrus.

**Figure 6.**
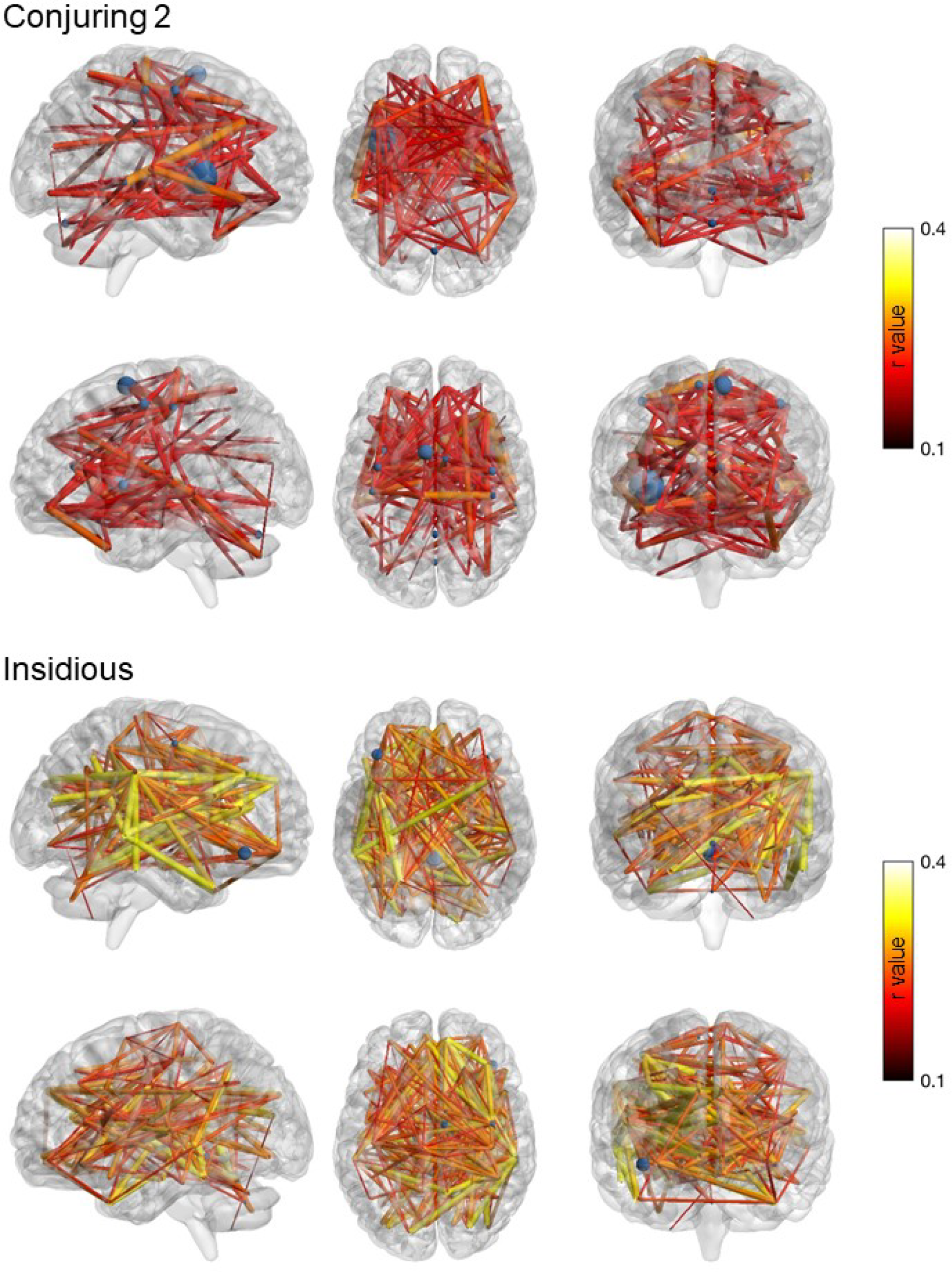
Seed-Based Phase Synchronization for each movie. Conjuring 2 (top, FDR corrected p = 0.05) and Insidious (bottom, FDR corrected p = 0.01). Phase similarity at each time point was calculated for each of 166 ROI pairs taken from the AAL atlas and correlated with the fear ratings. Connectome graphs (BrainNet Viewer: Xia, Wang, & He, 2013) depict those region pairs that exhibited a significant relationship between phase similarity and fear ratings with node size reflecting the number of connections and edge size and color reflecting the strength of the correlation. See **Supplementary Figure 3** for a depiction of these results as a correlation matrix.

Despite the consistent effects across both movies, some differences were also observed. The Conjuring 2 exhibited increased ISC as fear increased in the left cuneus, bilateral superior parietal lobe and, in the cerebellum, the cerebellar lingual and tonsil and uvula. The Conjuring 2 also exhibited several regions in the occipital lobe that showed a decrease in the ISC as fear increased (bilateral middle occipital gyri and bilateral lingual gyri, FDR corrected p = .05). Insidious exhibited additional fear related ISC in frontal regions (bilateral anterior medial frontal gyrus and middle frontal gyri), temporal regions (left anterior superior temporal gyrus, bilateral posterior superior temporal sulcus and right uncus), and occipital regions (bilateral superior occipital gyri). No negative associations were observed for Insidious.

### Fear-dependent changes in functional connectivity (Seed based Phase Synchronization)

At an FDR corrected alpha level of .05 (but not .01), fear predicted widespread functional connectivity across the brain for both movies, and several regions exhibited consistently high number of connections. These connectivity changes were also consistent across movies: the mean ICC coefficient for the unthresholded connectivity matrices across the two movies was 0.95 (indicative of excellent reliability, Koo & Li, 2016). Fear related connectivity within the frontal cortex was sparse, except for the precentral gyri, which acted as a hub for fear relevant connectivity with many areas within the frontal cortex. The right frontal middle gyrus exhibited increased functional connectivity with many regions in the occipital cortex, as well as the temporal and parietal cortices. The left postcentral gyrus increased fear related functional connectivity with several regions in the frontal lobe, and the cingulate cortex, which itself acted as a hub increasing connectivity with the temporal lobe and limbic system as fear increased. Fear predicted functional connectivity between left paracentral lobule and several regions in the frontal lobe, cingulate cortex, and posterior central gyrus. There were additional highly connected regions for each individual movie. The Conjuring 2 elicited increased connectivity as fear increased between the right frontal inferior orbital gyrus and the temporal lobe, of which the left superior temporal gyrus connected richly with the limbic system, whilst the left middle temporal lobe connected richly with the frontal lobe. The left middle frontal orbital gyrus in turn exhibited multiple connections with the temporal lobe. For Insidious, several frontal gyri (right inferior operculum, right inferior triangularis, right superior medial, left inferior orbital) exhibited fear associated connectivity with occipital and parietal regions, subcortical regions, and the cerebellum. Bilateral insula cortices increased connectivity with frontal and cingulate cortices, as well as the cerebellum. Bilateral transverse temporal gyri connected richly with anterior cingulate cortex, whilst in the partial lobe, the right supramarginal gyrus, right inferior parietal lobe, and bilateral postcentral gyri increased connectivity with frontal, cingulate, and occipital regions.

## Discussion

Our main finding was that acute threat elicited consistent activity in a distributed set of cortical, limbic, and cerebellar regions, most notably the prefrontal cortex, paracentral lobule, amygdala, cingulate cortex, insula, PAG, parrahippocampus, and thalamus. These regions have been previously identified as being active in response to threat (Mobbs et al., 2007; Qi et al, 2018; Zhu & Thagard, 2002). However, the activity of these regions was not associated with slower-frequency experience of fear, despite the high pass filter of 256s optimizing the GLM analysis to detect the low energy changes in fear ratings, which peaked at around 0.01Hz. Instead, these feelings of suspense were associated with increased activity in the sensory (both auditory and visual) cortices and a small portion of the parietal lobule. These differential patterns of activity suggest separable mechanisms for anticipation of threat, requiring increased perceptual and attentional focus, and acute responses to threat onset, requiring instinctive emotional processing centers, learning/memory, and action planning processes. Importantly, these effects were replicable in two independent samples of subjects and with two different stimulus movies, highlighting the replicability and generalizability of the results.

### Different timescales of fear

As fear becomes more imminent, amplified sensory processing and vigilance promote evidence gathering, whilst motor preparation (evidenced by activation in the precentral gyrus) promotes rapid protective responses whenever needed. The sudden onset of threat in turn elicits an abrupt increase in activity in regions associated with emotional processing, notably those processing the saliency of a stimulus (amygdala, Liberzon, Phan, Decker, & Taylor, 2003), and those triggering hyper-arousal required to act swiftly (PGA, Satpute et al., 2013) and avoid that which has previously been learnt to be harmful (hippocampus, Phelps, 2004). The thalamus may act as a relay between these areas and cortical regions involved in the homeostatic maintenance and emotional experience (insula cortex, Critchley, 2005; Phan, Wager, Taylor, & Liberzon, 2002), the formulation of motor plans to mitigate danger (anterior and posterior cingulate cortices, Beckmann, Johansen-Berg, & Rushworth, 2009), and the preparation of concrete motor acts in the precentral gyrus.

Interestingly, brain regions associated with the onset of acute fearful stimuli, despite not being associated with increasing fear at the individual level, did exhibit significant fear-dependent intersubject synchronization. That is, during high-fear episodes, brain activity became time-locked across subjects in several brain regions associated with the rapid onset of threat, notably those involved in the instigation of a rapid stress response (PGA) and the preparation and implementation of action (cingulate cortex, paracentral lobule). It is thus possible that sustained fear induces a reliable time-locked fluctuation in these regions, possibly reflecting the role that a commonly experienced increase in sustained fear has in the preparation of a predictable but highly constrained range of possible responses (freeze, fight, or flee). Importantly, this increased similarity at the neural level was also associated with increased similarity in subjective experience of fear: The more afraid the participants felt, the more similar their subjective feeling time courses became. This parallels with behavioral work showing that negative emotional states are associated with narrowing of mental focus and cognitive processing styles (Bishop, 2007; Panksepp, 1998).

### Functional networks for the fear response

Functional connectivity analysis revealed that, despite regional responses to sustained fear being modest, fear was associated with profound functional connectivity changes. This functional connectivity increased as fear increased, as if the cognitive anticipatory fear mechanism prepared the reactionary fear mechanism as threat became closer in spatiotemporal proximity. The frontal cortex housing complex threat appraisal and decision making mechanisms (Kalisch & Gerlicher, 2014; Rushworth, Buckley, Behrens, Walton, & Bannerman, 2007) exhibited increased connectivity with not only visual processing areas of the occipital cortex, but emotional processing areas of the limbic system. The frontal lobe and cingulate cortex (Etkin, Egner, & Kalisch, 2011) both exhibited fear related connectivity with the pre and post central gyri, possibly preparing concrete motor plans and escape behaviors in primary and supplementary motor cortices. This suggests that, although the anticipatory and reactionary fear networks may be dissociable in terms of absolute neural activity, they exhibit information transfer during threatening situations, whereby the sensory processing areas monitor for threat and resolve ambiguity, and engage emotional appraisal and action planning mechanisms as threat becomes more likely and the need for immediate action becomes more prescient (Fanselow, 1994; Lang, Davis, & Öhman, 2000; Lehne & Koelsch, 2015).

We propose that the anticipatory fear network weighs the emotional or threat related context of sensory information, and primes the reaction response network when threat subsequently arises. However, although sustained fear precipitated widespread connectivity throughout the brain, the effect of fear on the synchronization of this connectivity across individuals (see **Supplementary Analysis 2**) was far more limited and restricted to activity between regions implicated in the anticipation and response to threat itself. That is, the connectivity became increasingly time locked between regions associated with visual and emotional processing, and action planning and preparation. This suggests that fear not only synchronizes neural activity in these regions across individuals, but also the information transfer between these regions as the preparation to respond becomes increasingly similar.

Altogether our results establish that pre-encounter/anticipatory and post-encounter/reactionary networks do not work in isolation that require a qualitative shift depending on a discrete threshold of threat proximity. Instead, they work in concert throughout threat evaluation that gradually shifts from one to the other as threat increases in proximity. This insight would not have been possible using conventional model based approaches that require controlled stimuli discretely categorized into anticipatory and reactionary stimuli, and which would inevitably lead to the description of a binary system of pre and post threat onset (e.g., Mobbs et al., 2007; Qi et al., 2018). Instead, using naturalistic stimuli and a data driven approach that permits the reliability of neural activity to be established whilst accommodating the complex and dynamic nature of the neural signal and the realistic stimuli that elicits it (Glerean et al., 2012; Hasson et al., 2004), we were nevertheless able to not only confirm these systems, but reveal how they functionally interact during a fearful situation.

### Synchronous brain activation and contagion of fear

Emotion-dependent time-locking of brain activity across individuals could also provide the basis for transferring fear from individual to individual (Nummenmaa et al., 2018). Observing someone being afraid activates similar cortical and limbic regions as when experiencing the emotion oneself (de Gelder, Snyder, Greve, Gerard, & Hadjikhani, 2004; Nummenmaa, Hirvonen, Parkkola, & Hietanen, 2008) and induces functional connectivity between key areas such as the amygdala, anterior cingulate cortex, and anterior insula cortex (Yoshihara et al., 2016), suggesting that remapping of others’ emotional states in the limbic circuits could promote understanding others’ emotional states and response coordination. Predicting, understanding, and responding to the emotions of other people is crucial in social interactions, and is especially important with respect to fear. Other’s fearful reactions act as a cue to potential threats to ourselves (Tipples, 2006), and children learn what is dangerous by observing the fearful reactions of other people (Gerull & Rapee, 2002; Olsson & Phelps, 2007). Fearful faces attract attention (Bannerman, Temminck, & Sahraie, 2012), and simply seeing someone express fear elicits a sympathetic fearful response (Nummenmaa, Glerean, et al., 2014; Vaughan & Lanzetta, 1980; Yoshihara et al., 2016). Fear is alleviated when in the company of others (Hennessy, Kaiser, & Sachser, 2009), and social bonding is fostered after sharing a traumatic experience (Bastian, Jetten, & Ferris, 2014; Jong, Whitehouse, Kavanagh, & Lane, 2015). Because behavioral and neural synchronization across individuals is consistently associated with social bonding (see review in Nummenmaa, Lahnakoski, & Glerean, 2018), it is possible that the fear-triggered synchronous brain activity and behavior in a group could also promote their social bonding.

### Conclusions

Our combination of model-based and model-free approaches for naturalistic neuroimaging data reveals dynamic interaction between two separable systems for the anticipation of threat from environmental cues, and the reaction to threat onset, with a temporal shift between them as the spatiotemporal proximity of threat decreases, and anticipatory planning mechanisms inform subsequent responses. These effects are reliable across subjects and experimental conditions, further highlighting the feasibility of naturalistic stimulation models in understanding brain basis of emotions.

## SUPPLEMENTARY MATERIAL

**Supplementary Figure 1.**
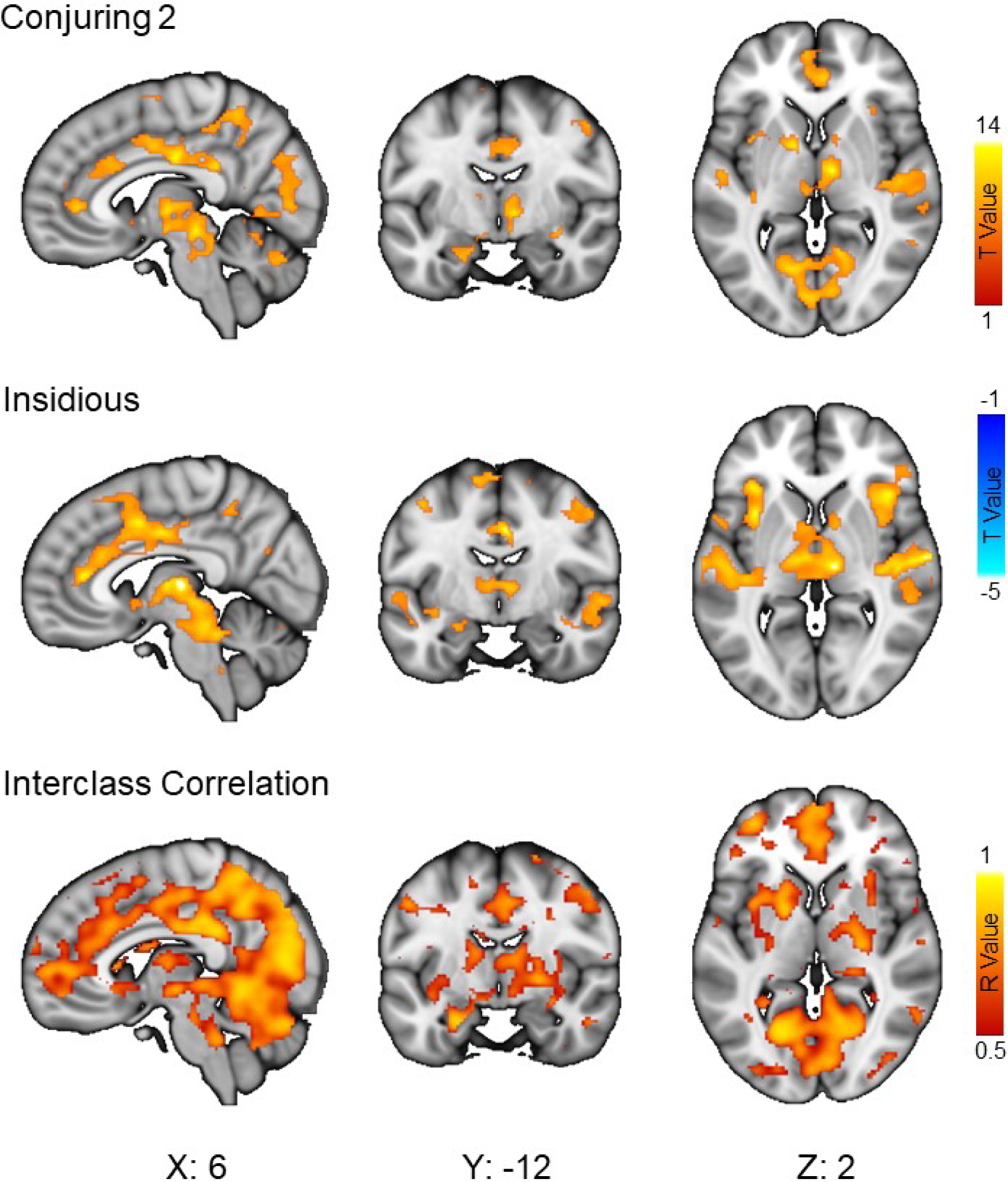
The effect of Jump-scares on neural activity for each movie (Top: Conjuring 2; FDR corrected *p* = .05; Middle: Insidious, uncorrected *p* = .001) and the interclass correlation coefficient map for the two movies (bottom). Jump-scare onsets were modelled with a stick function using general linear model analysis. The ICC (one-way random effects between-subjects for each voxel of the individual t contrast maps) was thresholded at *r* = 0.5 indicative of moderate or more reliability (Koo & Li, 2016).

**Supplementary Figure 2.**
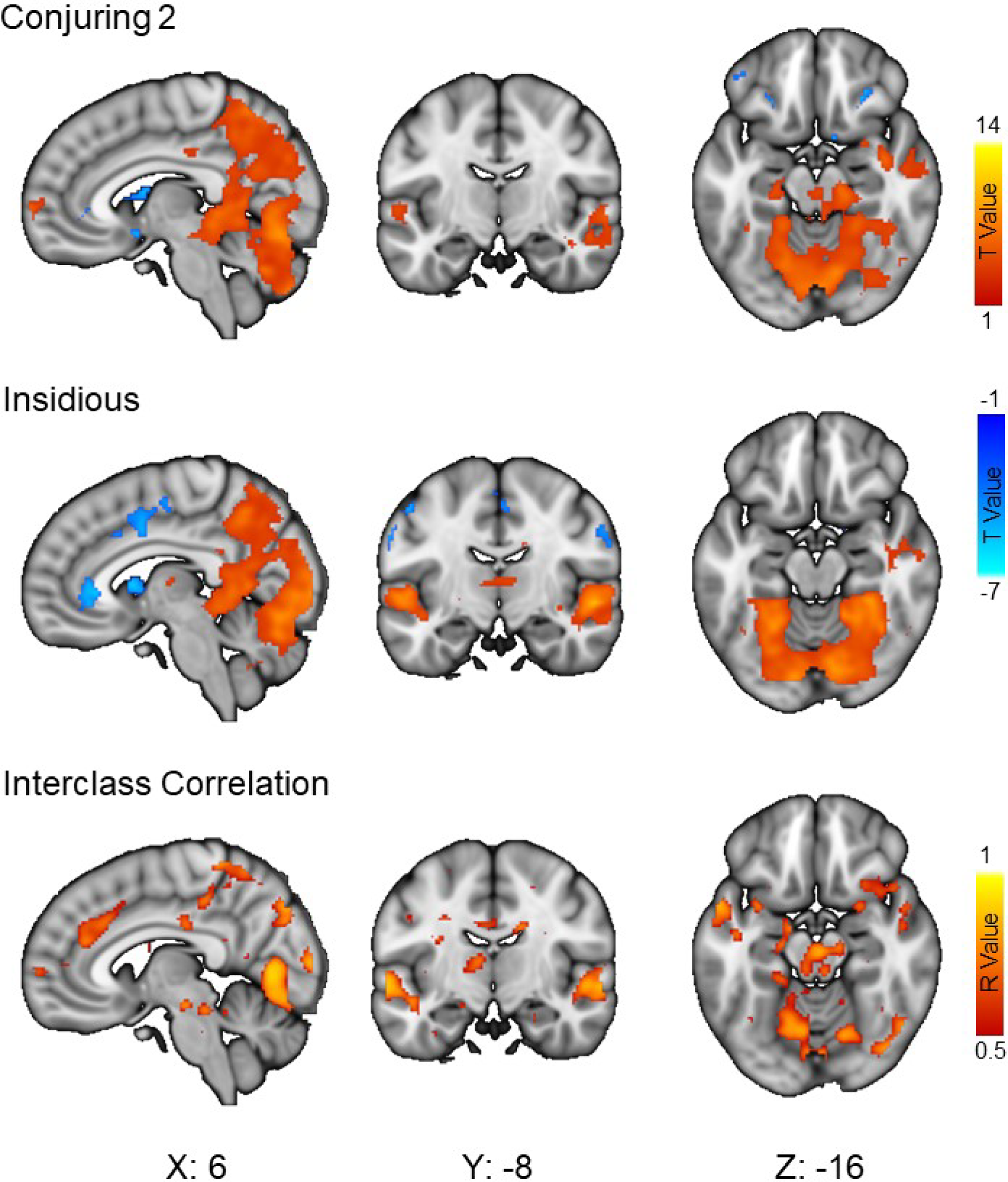
The relationship between experienced fear and neural activity for each movie (Top: Conjuring 2; Middle: Insidious) and the interclass correlation coefficient map for the two movies (bottom). Fear ratings were convolved with a hemodynamic response function and entered as a regressor into a GLM analysis with a high-pass band filter of 256s (uncorrected *p* = 0.001). The ICC (one-way random effects between-subjects for each voxel of the individual t contrast maps) was thresholded at *r* = 0.5 indicative of moderate or more reliability (Koo & Li, 2016).

**Supplementary Figure 3.**
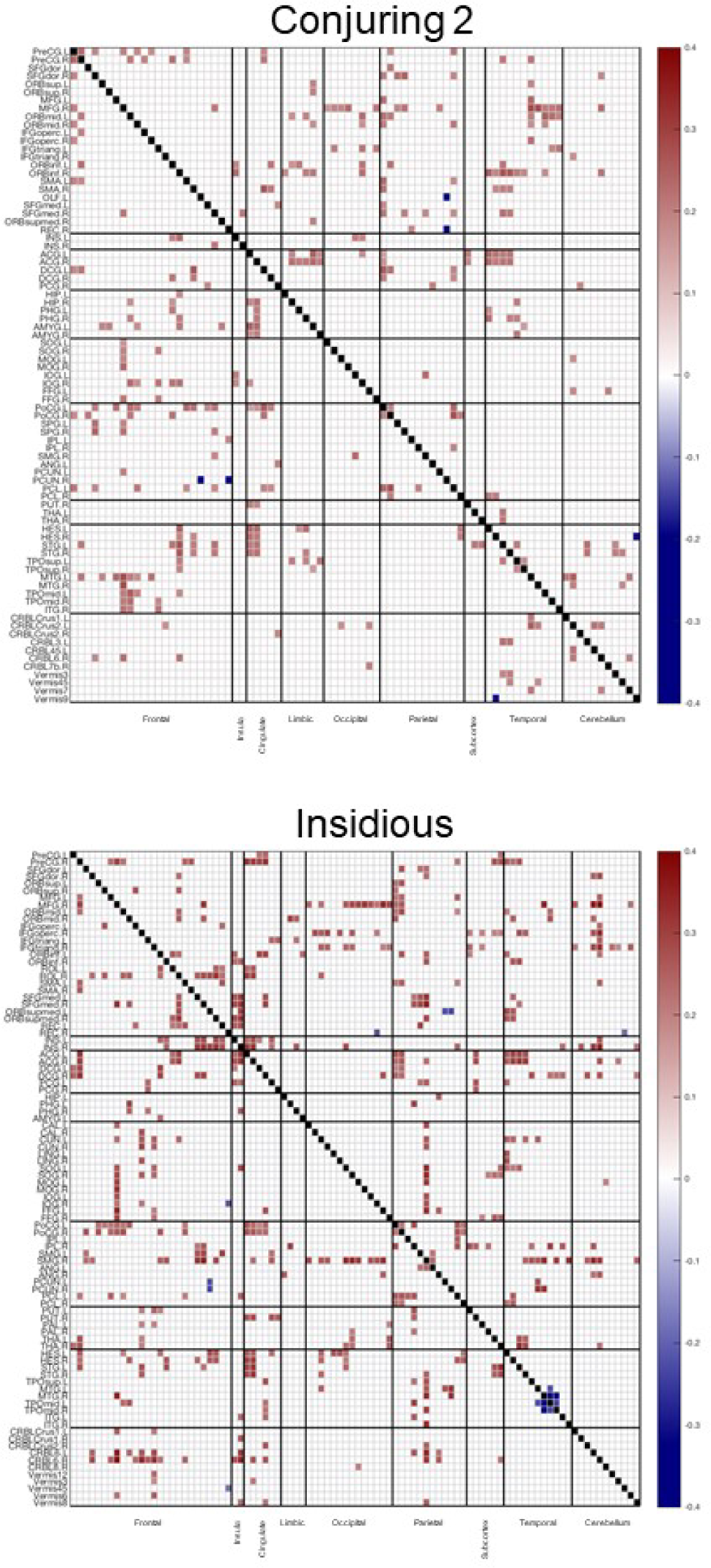
Coefficient matrices of the relationship between fea and seed-based phase synchronization for each movie: Conjuring 2 (top, FDR corrected p = 0.05) and Insidious (bottom, FDR corrected p = 0.01). Phase similarity at each time point was calculated for each of 166 ROI pairs taken from the AAL atlas and correlated with the fear ratings. Correlations matrices depict those node pairs that exhibited a significant correlation between phase similarity and fear ratings. The mean ICC coefficient for the unthresholded correlation matrices across the two movies was 0.97 (indicative of excellent reliability, Koo & Li, 2016).

### Supplementary Analysis 1

#### Fear induced changes in intersubject synchronization of functional connectivity

Inter-subject Seed Based Phase Synchronization (ISBPS, https://github.com/eglerean/funpsy, Glerean et al., 2012) was employed to investigate how functional connectivity between brain regions is synchronized between individuals, and how this is influenced by emotion. ISBPS reveals the extent to which functional connectivity between any given region pair is synchronized between individuals. That is, not only is there correlated activity between regions, but this is time locked across individuals, suggesting that the connectivity is the same for each individual at any given moment. When correlated with the fear ratings, this reveals how the similarity in functional connectivity across individuals increases with rising intensities of emotion.

The phase similarity between each region pair within a given participant is subject to an inter-subject phase synchronization whereby a full subject-by-subject phase difference is computed. This provides a dynamic measure of the reliability of functional connectivity across individuals, and enables us to establish not only whether functional connectivity occurs for each individual, but how synchronized this functional connectivity is over time. This was then correlated with the fear ratings to demonstrate how fear affects the synchronization of connectivity between regions across the whole group. These correlations were conducted with an autocorrelated degrees of freedom (Conjuring 2: 150 to 649; Insidious: 123 to 541) with an FDR corrected p value of 0.001.

##### Conjuring 2

Connectivity between many regions within the frontal cortex exhibited increased synchronization between individuals as fear increased. Notably, the precentral gyri acted as a hub not only within the frontal lobe, but also with the insula cortex, parietal lobe, and thalamus. The insula exhibited functional connectivity synchronization as fear increased with regions throughout the frontal lobe, cingulate cortex, parietal lobe, limbic system, and cerebellum. The left supramarginal gyrus also exhibited synchronized functional connectivity as fear increased with many regions throughout frontal and parietal lobes, cingulate and insulate cortices, and cerebellum. Further findings of note were a fear related synchronization of connectivity between the anterior cingulate cortex and precentral gyrus, and between the left thalamus and several frontal areas.

##### Insidious

Bilateral thalamus exhibited increased synchronization of functional connectivity with several frontal regions, the right insula, bilateral cingulate gyri, right hippocampus and culmen of the cerebellum. The culmen in turn showed extensive synchronized connectivity as fear increased with the limbic system, left paracentral lobule, right middle cingulate cortex, bilateral insula, and several frontal regions. Fear was also a predictor of synchronized connectivity between the right inferior frontal orbital cortex and the culmen, right amygdala and right hippocampus, and between bilateral precentral gyri, and right middle cingulate cortex.

**Supplementary Figure 4.**
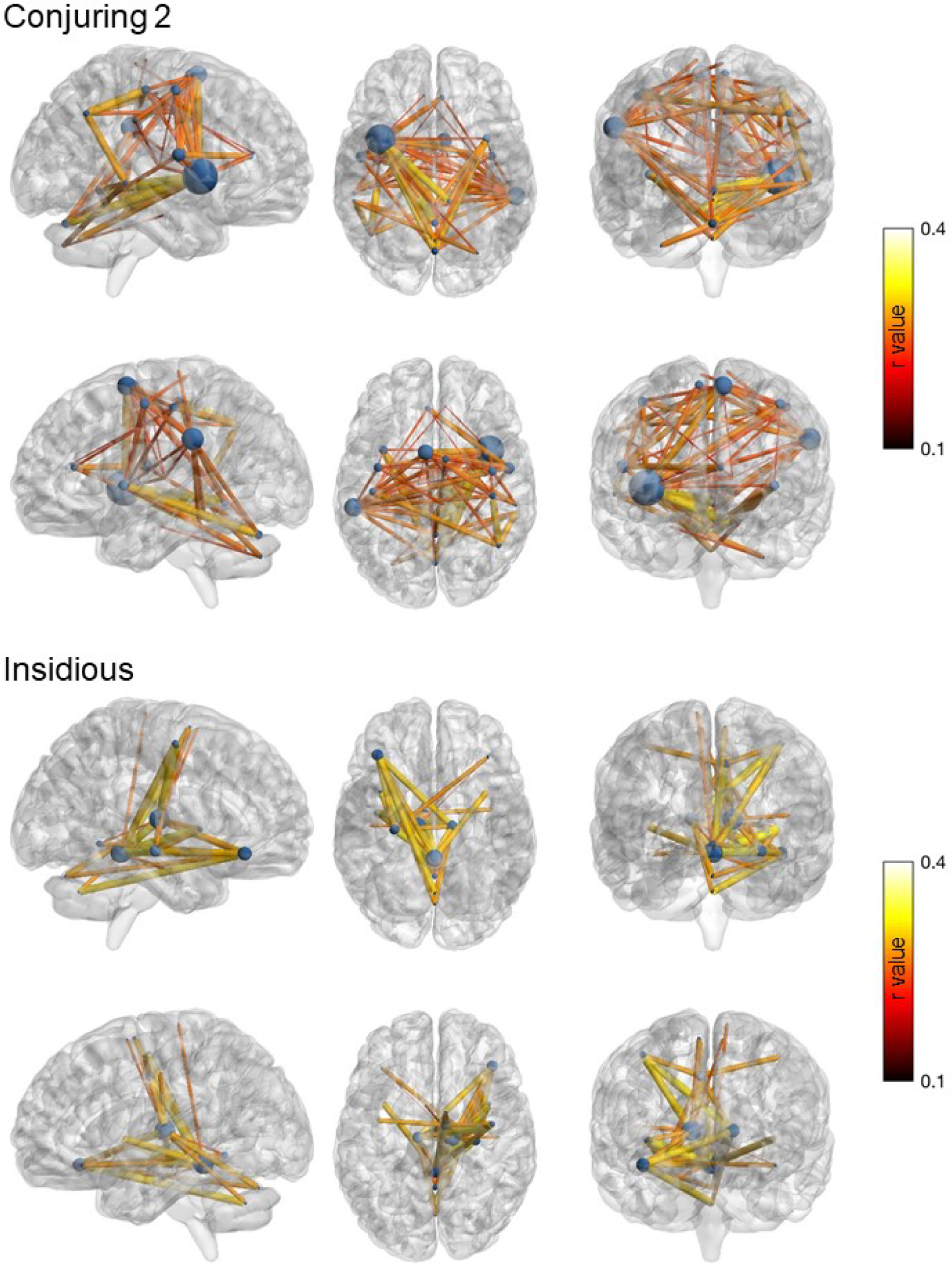
The relationship between fear and intersubject synchronization of functional connectivity for Conjuring 2 (top) and Insidious (bottom) (both FDR corrected p = 0.001). Phase similarity at each time point between each region pair was subject to inter-subject phase similarity analysis, and correlated with the fear ratings. Connectome graphs (BrainNet Viewer: Xia, Wang, & He, 2013) depict those region pairs whose phase similarity exhibited increased intersubject synchronization as fear increased, with node size reflecting the number of connections and edge size and color reflecting the strength of the correlation. See **Supplementary Figure 5** for a depiction of these results as a correlation matrix.

**Supplementary Figure 5.**
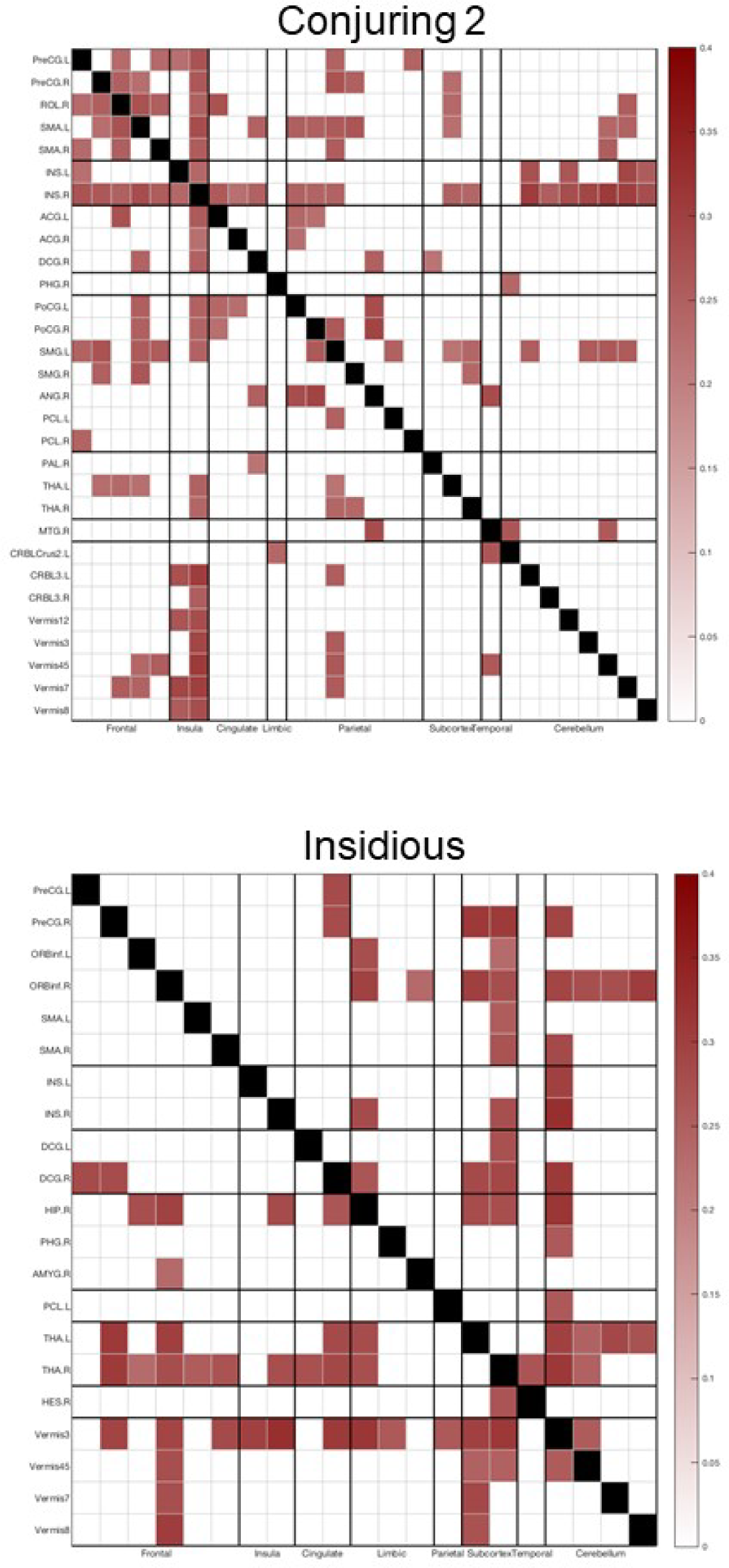
Intersubject Seed Based Phase Synchronization for Conjuring 2 (top) and Insidious (bottom) (both FDR corrected p = 0.001). Phase similarity at each time point between each region pair was subject to inter-subject phase similarity analysis, and correlated with the fear ratings. Correlations matrices depict those node pairs that exhibited a significant correlation between phase similarity and fear ratings. The mean ICC coefficient for the unthresholded correlation matrices across the two movies was 0.95 (indicative of excellent reliability, Koo & Li, 2016).

### Supplementary Analysis 2

#### Horror Movie Survey

Participants (N = 216, 62% Male, 37% Female, mean age = 29.1 years) were recruited online via snowballing sampling through university networks, and by targeting online horror movie chat rooms. Respondents were predominantly from Finland (76.4%), but also the USA (8.8%), Canada (2.3%), United Kingdom (1.9%), amongst others (Australia, Belgium 1.4% each, France, Germany, Greece, Ireland, Sri Lanka, Thailand 0.9% each, Estonia, Lithuania, Poland, Turkey 0.5% each).

Participants first completed a few questions about their movie going habits in general, and specifically with respect to horror movies. The sample were regular and frequent movie goers, with three quarters of respondents reporting watching a movie at least once every 2 weeks, with the majority of these watching a movie on a weekly basis. Almost half of the sample reported that they liked horror movies, and many reported watching a horror movie once a month or once every 3 months. Only 10% reported never watching a horror movie. When asked how horror movies made them feel, the most frequent responses were predominantly feelings of high arousal and negative valence associated with fear (scared, anxious, nervous, confused), but also with positive valence feelings of excitation and pleasure. Negative feelings associated with the expulsion of noxious substances, such as disgust and nausea, were also common. Participants reported a wide variety of different sources of horror to be the scariest, but psychological fear associated with manipulating the mind of either the viewer and/or the protagonist was the most common. Movies that instilled fear with either extreme realism or extreme unreal supernatural elements were also the most likely to elicit fear. In support of the contention that ambiguity and uncertainty elicit fear, the majority of participants stated that the idea of something that wasn’t on screen was far scarier than actually seeing something. Lastly, over half of respondents reported that they preferred watching horror movies with others, which suggests that horror movies are, like movie watching in general, a very social experience. However, the uniquely intense emotions elicited by horror movies require the comfort that other’s company provides, and the social bonding that this promotes may provide one of the key psychological functions of horror movies.

Participants were then provided with a list of the 100 greatest horror movies of the past 100 years and asked if they had seen them and, if so, how good and how scary it was on a scale of 1 to 10. We had no a priori hypotheses for this data, but an exploratory correlation analysis with the variables year of production, scariness ratings, and quality ratings (Bonferroni corrected alpha = 0.017) revealed that scariness and quality were positively correlated (*r* = .528, *p* < .001), and whilst scariness was marginally correlated with year of production (*r* = .229, *p* = .02), there was no indication that movie quality increased with more recent production years (*r* = -.02, *p* = .843).

**Q1. How often do you watch a movie? Once…**

**Q1. How often do you watch a movie? Once...**
| a day | a week | 2 weeks | a month | 3 months | 6 month | a year |
| --- | --- | --- | --- | --- | --- | --- |
| 13.0% | 40.3% | 22.2% | 15.7% | 7.4% | 0.9% | 0.5% |

**Q2. Do you like horror movies?**

| No | Sometimes | Yes |
| --- | --- | --- |
| 18.1% | 31.5% | 47.7% |

**Q3. How often do you watch a horror movie? Once…**

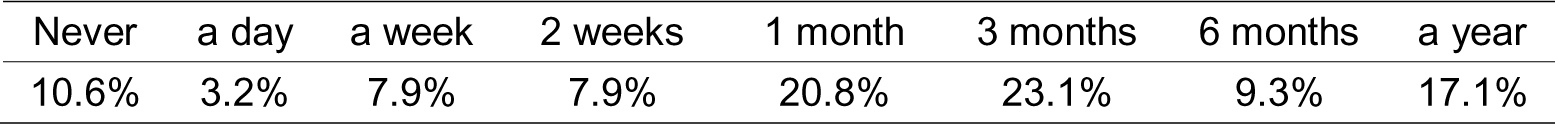

**Q4. What percentage of movies that you watch are horror movies?**

| 0 | 10 | 20 | 30 | 40 | 50 | 60 | 70 | 80 | 90 |
| --- | --- | --- | --- | --- | --- | --- | --- | --- | --- |
| 18.9% | 25.9% | 17.1% | 10.6% | 5.1% | 7.9% | 4.6% | 4.2% | 4.6% | 0.9% |

**Q5. How do horror movies make you feel? (tick all that apply)**

| Feeling | Count |
| --- | --- |
| Excited | 152 |
| Scared | 138 |
| Anxiety | 122 |
| Nervous | 104 |
| Disgusted | 69 |
| Pleasure | 61 |
| Confused | 52 |
| Nauseated | 29 |
| Frustrated | 3 |
| Relieved | 3 |
| Bored | 2 |
| Hilarity | 2 |
| Insecurity | 2 |
| Wonder | 2 |
| Courageous | 1 |
| Curiosity | 1 |
| Escapism | 1 |
| Intrigued | 1 |
| Joy | 1 |
| Numb | 1 |
| Paranoid | 1 |
| Ridicule | 1 |
| Surprised | 1 |
| Thrill | 1 |
| Uncomfortable | 1 |

**Q6. What type of horror movie is the scariest? (tick all that apply)**

| Type | Count |
| --- | --- |
| Psychological horror | 161 |
| Real events | 130 |
| Supernatural | 112 |
| Human criminals | 79 |
| Torture | 75 |
| Blood and gore | 41 |
| Survival horror | 38 |
| Slasher | 32 |
| Monsters | 24 |
| Sci-fi horror | 13 |
| Gothic horror | 7 |
| Teen horror | 5 |

**Q7. Which is scarier?**

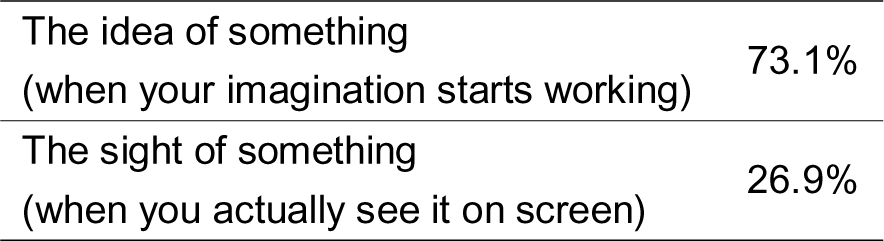

|  |  |
| --- | --- |
| The idea of something<br>(when your imagination starts working) | 73.1% |
| The sight of something<br>(when you actually see it on screen) | 26.9% |

**Q8. Do you prefer to watch horror movies…**

**Q8. Do you prefer to watch horror movies...**
|  |  |
| --- | --- |
| Alone | 11.1% |
| It doesn't matter | 32.4% |
| With other people | 56.5% |

**Supplementary Table 1.** Full results of the horror movie survey, indicating the percentage of respondents who had seen the movie, and their ratings of the movie’s scariness and quality. The number of jump scares are also indexed (where available). Movies are ranked according to scariness.

Horror Movie Ratings
| Title | Year | Have you seen this movie? |  |  |  |  |  |
| --- | --- | --- | --- | --- | --- | --- | --- |
|  |  | Yes | No, but heard of it | Not seen nor heard of it | Scary | Quality | Jump scares |
| The Devil's Backbone | 2001 | 4.2 | 6.3 | 89.6 | 8.5 | 10.0 | 4 |
| The Wailing | 2016 | 2.1 | 8.3 | 89.6 | 8.0 | 9.0 | 4 |
| The Conjuring | 2013 | 39.6 | 31.3 | 29.2 | 7.5 | 7.5 | 12 |
| REC 2 | 2009 | 10.4 | 29.2 | 60.4 | 7.3 | 5.8 | - |
| **Insidious** | **2010** | **29.2** | **25.6** | **45.2** | **7.1** | **7.1** | **24** |
| The Exorcist | 1973 | 53.0 | 23.2 | 23.8 | 7.1 | 7.0 | 10 |
| Goodnight Mommy | 2015 | 4.2 | 10.4 | 85.4 | 7.0 | 8.0 | 1 |
| A Chinese Ghost Story | 1987 | 1.8 | 7.1 | 91.1 | 7.0 | 7.8 | - |
| **The Conjuring 2** | **2016** | **37.5** | **31.0** | **31.5** | **7.0** | **7.3** | **22** |
| Under The Shadow | 2016 | 2.1 | 14.6 | 83.3 | 7.0 | 7.0 | 9 |
| Sinister | 2012 | 35.1 | 30.4 | 34.5 | 7.0 | 6.8 | 17 |
| The Ring | 2002 | 61.9 | 31.5 | 6.5 | 7.0 | 6.5 | 9 |
| The Shining | 1980 | 81.3 | 12.5 | 6.3 | 6.8 | 7.8 | 3 |
| The Night Of The Hunter | 1955 | 4.2 | 10.1 | 85.7 | 6.7 | 9.0 | - |
| We Are What We Are | 2013 | 6.3 | 8.3 | 85.4 | 6.5 | 6.8 | - |
| Paranormal Activity | 2007 | 51.8 | 35.1 | 13.1 | 6.5 | 5.9 | 10 |
| The Orphanage | 2007 | 33.3 | 33.3 | 33.3 | 6.4 | 6.8 | 6 |
| The Silence Of The Lambs | 1991 | 54.2 | 31.5 | 14.3 | 6.4 | 8.6 | 0 |
| A Nightmare On Elm Street | 1984 | 47.9 | 37.5 | 14.6 | 6.4 | 6.6 | 11 |
| The Descent | 2005 | 14.9 | 19.0 | 66.1 | 6.2 | 6.8 | 16 |
| The Others | 2001 | 27.4 | 25.6 | 47.0 | 6.2 | 7.8 | 4 |
| Poltergeist | 1982 | 37.5 | 47.9 | 14.6 | 6.1 | 6.2 | 9 |
| 28 Days Later... | 2002 | 27.4 | 25.6 | 47.0 | 6.1 | 6.4 | 10 |
| Slither | 2006 | 13.1 | 34.5 | 52.4 | 6.1 | 4.9 | 10 |
| Alien | 1979 | 43.8 | 35.4 | 20.8 | 6.1 | 7.7 | 11 |
| Train To Busan (Bu-San-Haeng) | 2016 | 3.0 | 20.2 | 76.8 | 6.0 | 7.0 | 4 |
| Halloween | 1978 | 37.5 | 50.0 | 12.5 | 6.0 | 6.2 | 13 |
| The Innocents | 1961 | 2.1 | 22.9 | 75.0 | 6.0 | 6.0 | 4 |
| The Witch | 2016 | 10.1 | 16.1 | 73.8 | 5.9 | 6.9 | 4 |
| It Follows | 2014 | 29.2 | 20.8 | 50.0 | 5.9 | 6.7 | 5 |
| Hush | 2016 | 29.8 | 24.4 | 45.8 | 5.8 | 6.5 | 7 |
| Rosemary's Baby | 1968 | 26.2 | 27.4 | 46.4 | 5.8 | 6.7 | 0 |
| The Texas Chainsaw Massacre | 1974 | 35.1 | 48.2 | 16.7 | 5.7 | 5.8 | 2 |
| The Woman In Black | 2012 | 22.9 | 27.1 | 50.0 | 5.7 | 6.4 | 16 |
| Don't Breathe | 2016 | 15.5 | 21.4 | 63.1 | 5.7 | 7.1 | 17 |
| The Babadook | 2014 | 33.3 | 25.0 | 41.7 | 5.7 | 5.9 | 11 |
| The Ring 2 | 2005 | 35.4 | 56.3 | 8.3 | 5.6 | 5.0 | 11 |
| Lights Out | 2016 | 20.8 | 29.2 | 50.0 | 5.6 | 5.8 | 19 |
| Misery | 1990 | 20.8 | 16.7 | 62.5 | 5.6 | 7.7 | 3 |
| Psycho | 1960 | 45.8 | 37.5 | 16.7 | 5.6 | 7.6 | 2 |

| Have you seen this movie? |  |  |  |  |  |  |  |
| --- | --- | --- | --- | --- | --- | --- | --- |
| Title | Year | Yes | No, but<br>heard of it | Not seen nor<br>heard of it | Scary | Quality | Jump<br>scores |
| The Vanishing (Spoorloos) | 1988 | 3.0 | 20.2 | 76.8 | 5.6 | 6.5 | - |
| The House Of The Devil | 2009 | 10.4 | 16.7 | 72.9 | 5.6 | 6.2 | 4 |
| Aliens | 1986 | 28.0 | 42.3 | 29.8 | 5.5 | 6.2 | 13 |
| Pontypool | 2008 | 2.4 | 8.3 | 89.3 | 5.5 | 6.3 | 1 |
| 10 Cloverfield Lane | 2016 | 17.3 | 23.2 | 59.5 | 5.4 | 5.9 | 8 |
| Dead Of Night | 1945 | 6.3 | 12.5 | 81.3 | 5.3 | 7.7 | - |
| Shutter | 2004 | 12.5 | 26.2 | 61.3 | 5.3 | 5.5 | 11 |
| The Haunting | 1963 | 8.3 | 29.2 | 62.5 | 5.3 | 5.3 | 3 |
| The Fly | 1986 | 33.3 | 22.9 | 43.8 | 5.2 | 6.4 | 2 |
| Invasion Of The Body Snatchers | 1956 | 9.5 | 22.0 | 68.5 | 5.1 | 6.3 | - |
| Night Of The Living Dead | 1968 | 17.3 | 41.7 | 41.1 | 5.1 | 6.3 | - |
| The Mist | 2007 | 37.5 | 25.0 | 37.5 | 5.1 | 6.7 | 5 |
| Repulsion | 1965 | 2.1 | 8.3 | 89.6 | 5.0 | 7.0 | 7 |
| Eyes Without A Face | 1960 | 4.2 | 16.7 | 79.2 | 5.0 | 9.0 | - |
| Green Room | 2016 | 6.3 | 22.9 | 70.8 | 5.0 | 7.5 | 1 |
| Don't Look Now | 1973 | 7.1 | 4.2 | 88.7 | 4.9 | 6.5 | - |
| Jaws | 1975 | 66.7 | 25.0 | 8.3 | 4.7 | 6.1 | 4 |
| Henry: Portrait Of A Serial Killer | 1986 | 8.3 | 22.9 | 68.8 | 4.7 | 7.0 | - |
| Scream | 1996 | 64.6 | 31.3 | 4.2 | 4.7 | 5.0 | 19 |
| Shin Godzilla | 2016 | 3.0 | 36.3 | 60.7 | 4.6 | 6.3 | - |
| I Walked With A Zombie | 1943 | 1.2 | 7.1 | 91.7 | 4.5 | 6.0 | - |
| The Cabin In The Woods | 2012 | 41.7 | 35.4 | 22.9 | 4.5 | 6.6 | 13 |
| The Girl With All The Gifts | 2016 | 3.0 | 7.1 | 89.9 | 4.4 | 7.8 | 3 |
| Carrie | 1976 | 38.1 | 33.3 | 28.6 | 4.4 | 5.4 | 1 |
| The Cabinet Of Dr. Caligari | 1920 | 4.2 | 11.3 | 84.5 | 4.4 | 6.5 | - |
| What Ever Happened To Baby Jane? | 1962 | 7.7 | 26.2 | 66.1 | 4.3 | 6.6 | - |
| Pan's Labyrinth | 2006 | 36.9 | 21.4 | 41.7 | 4.3 | 6.8 | 6 |
| Drag Me To Hell | 2009 | 27.1 | 18.8 | 54.2 | 4.3 | 5.0 | 23 |
| Eraserhead | 1977 | 10.7 | 13.1 | 76.2 | 4.2 | 5.7 | - |
| Split | 2016 | 11.9 | 32.1 | 56.0 | 4.2 | 7.3 | 3 |
| An American Werewolf In London | 1981 | 13.7 | 25.0 | 61.3 | 4.1 | 5.5 | 14 |
| Evil Dead 2 | 1987 | 13.7 | 29.2 | 57.1 | 4.1 | 6.4 | 27 |
| Videodrome | 1982 | 5.4 | 13.1 | 81.5 | 4.0 | 6.3 | - |
| Bone Tomahawk | 2015 | 4.2 | 12.5 | 83.3 | 4.0 | 4.5 | 0 |
| A Field In England | 2014 | 2.1 | 8.3 | 89.6 | 4.0 | 9.0 | - |
| 28 Weeks Later... | 2007 | 14.6 | 31.3 | 54.2 | 4.0 | 6.6 | 17 |
| Cat People | 1942 | 3.6 | 16.1 | 80.4 | 3.9 | 7.5 | 2 |
| Let Me In | 2010 | 10.7 | 22.0 | 67.3 | 3.9 | 6.5 | 3 |
| World War Z | 2013 | 28.6 | 41.7 | 29.8 | 3.8 | 6.2 | 7 |
| Nosferatu The Vampire | 1922 | 12.5 | 52.1 | 35.4 | 3.8 | 6.0 | - |

**Supplementary Table 1.** Full results of the horror movie survey, indicating the percentage of respondents who had seen the movie, and their ratings of the movie's scariness and quality.
| Title | Year | Have you seen this movie? |  |  |  |  | Jump scores |
| --- | --- | --- | --- | --- | --- | --- | --- |
|  |  | Yes | No, but heard of it | Not seen nor heard of it | Scary | Quality |  |
| Predator | 1987 | 35.4 | 35.4 | 29.2 | 3.7 | 6.7 | 7 |
| I Am Legend | 2007 | 33.3 | 37.5 | 29.2 | 3.6 | 5.5 | 9 |
| Ginger Snaps | 2000 | 5.4 | 8.9 | 85.7 | 3.6 | 4.3 | - |
| Grindhouse | 2007 | 13.1 | 13.7 | 73.2 | 3.4 | 7.1 | - |
| Fright Night | 1985 | 16.7 | 27.1 | 56.3 | 3.4 | 5.1 | 10 |
| Let The Right One In | 2008 | 25.0 | 2.1 | 72.9 | 3.3 | 6.7 | 2 |
| Donnie Darko | 2004 | 31.3 | 14.6 | 54.2 | 3.3 | 7.5 | 3 |
| Freaks | 1932 | 3.0 | 13.1 | 83.9 | 3.3 | 7.8 | - |
| Re-Animator | 1985 | 6.0 | 12.5 | 81.5 | 3.0 | 5.5 | 4 |
| The Bride Of Frankenstein | 1935 | 8.3 | 54.2 | 37.5 | 3.0 | 4.7 | - |
| Frankenstein | 1931 | 20.8 | 69.0 | 10.1 | 2.9 | 5.6 | - |
| The Host | 2006 | 7.1 | 23.2 | 69.6 | 2.9 | 5.0 | - |
| Godzilla | 1956 | 16.7 | 68.8 | 14.6 | 2.9 | 5.0 | - |
| Phantom Of The Opera | 1925 | 22.9 | 60.4 | 16.7 | 2.8 | 6.7 | - |
| Dead Alive | 1992 | 5.4 | 19.6 | 75.0 | 2.7 | 6.2 | 3 |
| King Kong | 1933 | 26.8 | 58.3 | 14.9 | 2.6 | 5.1 | - |
| Cloverfield | 2008 | 25.0 | 22.9 | 52.1 | 2.3 | 5.5 | 7 |
| Attack The Block | 2011 | 10.4 | 4.2 | 85.4 | 2.3 | 5.4 | 12 |
| Under The Skin | 2013 | 4.2 | 16.7 | 79.2 | 2.0 | 2.5 | - |
| What We Do In The Shadows | 2014 | 5.4 | 10.7 | 83.9 | 1.0 | 6.0 | - |
| Dracula: Pages From A Virgin's Diary | 2003 | 2.1 | 27.1 | 70.8 | 1.0 | 6.0 | - |
| A Girl Walks Home Alone At Night | 2014 | 2.1 | 16.7 | 81.3 | 1.0 | 1.0 | 0 |
| Vampyr - Der Traum Des Allan Grey | 1932 | 2.1 | 10.4 | 87.5 | 1.0 | 6.0 | - |
| Backcountry | 2015 | 0.0 | 6.0 | 94.0 |  |  | 4 |

